# Genome-wide allele frequency changes reveal that dynamic metapopulations evolve differently

**DOI:** 10.1101/2024.01.18.575980

**Authors:** Pascal Angst, Christoph R. Haag, Frida Ben-Ami, Peter D. Fields, Dieter Ebert

## Abstract

**Background:** Two important characteristics of metapopulations are extinction– (re)colonization dynamics and gene flow between subpopulations. These processes can cause strong shifts in genome-wide allele frequencies that are generally not observed in “classical” (large, stable, panmictic) populations. Subpopulations founded by one or a few individuals, the so-called propagule model, are initially expected to show intermediate allele frequencies at polymorphic sites until natural selection and genetic drift drive allele frequencies toward a mutation-selection-drift equilibrium characterized by a negative exponential-like distribution of the site frequency spectrum (SFS).

**Results:** We followed changes in SFS distribution in a natural metapopulation of the cyclically parthenogenetic pond-dwelling microcrustacean *Daphnia magna* using biannual pool-seq samples collected over a five-year period from 118 ponds occupied by subpopulations of known age. As expected under the propagule model, SFSs in newly founded subpopulations trended toward intermediate allele frequencies and shifted toward right skewed distributions as the populations aged. Immigration and subsequent hybrid vigor altered this dynamic.

**Conclusions:** The analysis of SFS dynamics is a powerful approach to understand evolution in metapopulations. It allowed us to disentangle evolutionary processes occurring in a natural metapopulation, where many subpopulations evolve in parallel. Thereby, stochastic processes like founder and immigration events lead to a pattern of subpopulation divergence, while genetic drift, leads to converging SFS distributions in the persisting subpopulations. The observed processes are well explained by the propagule model and highlight that metapopulations evolve differently from classical populations.

## Background

In heterogeneous landscapes, populations are often fragmented, forming a patchwork of subpopulations connected by gene flow. Subpopulations may display strong dynamics like local extinction and habitat patch (re)colonization (1,2) that may create strong evolutionary changes in genome-wide allele frequencies, i.e., in the site frequency spectrum (SFS). The evolution of such metapopulations is, therefore, expected to differ from evolution in classical populations, here defined as large, stable, and panmictic populations, which are predominantly considered in population genetic studies (3). The SFS of classical populations is shaped by a combination of deterministic (e.g., natural selection) and stochastic (e.g., genetic drift) evolutionary processes (4). These processes also shape the SFSs of metapopulations, but their relative impact is expected to differ (5). In dynamic metapopulations, founder events, i.e., strong bottlenecks when a single or a few individuals colonize vacant habitat patches, are frequent and are expected to result in low genetic diversity and a higher abundance of alleles with intermediate frequencies (6,7). In the extreme case, genetic bottlenecks are caused by a single selfing colonizer, founding a subpopulation in which all polymorphic alleles are initially at frequencies close to 50 %. We therefore expect the SFSs of subpopulations in dynamic metapopulations to be skewed more toward intermediate frequencies than the right-skewed, negative exponential-like distributions observed in classical populations (8). However, this theoretical predication has not been assessed. Here, we quantify the distribution of genome-wide allele frequencies in a natural metapopulation and examine the processes that shape it over time.

The propagule model of colonization is often used to explain population genetic processes in metapopulations. It assumes that subpopulations are colonized by one or a few individuals, often originating from a single source population (9). After the founder bottleneck, evolutionary processes act on the intermediate frequency skewed distribution of the SFS (5). In the subpopulations, which are initially small and prone to inbreeding (high genetic load) (10,11), genetic drift dominates the evolutionary process so that the efficacy of deterministic processes like purifying or spatially varying selection is low (12,13). In just a few (sexual) generations genetic drift may reduce the skew of the SFS from intermediate-frequency alleles to a successively flattening SFS. Pervasive genetic drift and the occurrence of new mutations shift the SFS further toward mutation-drift equilibrium with a negative exponential-like distribution similar to those known from classical populations (8,14). However, this long-term effect is unlikely to contribute much to the observed SFS dynamics in dynamic metapopulations because subpopulations may not persist long enough. Gene flow is another important evolutionary process that shapes SFSs in metapopulations. It introduces new alleles into subpopulations, thus accelerating the evolution of SFSs toward a right skewed distribution (15). Gene flow may also lead to hybrid vigor, which is expected when residents and immigrant individuals mate to produce outbred offspring, especially if one or both parents suffered from high genetic load (11,16). Under hybrid vigor, the hybrid offspring has high fitness, i.e., a selective advantage over the resident population and the immigrant parent, because deleterious mutations, which are homozygous in the parents, are masked in a recessive state. Therefore, immigrant alleles can quickly increase in frequency, up to 50 % in the most extreme cases (11,16), leading SFSs to evolve toward alleles with intermediate frequencies and higher heterozygosity. Selection for heterozygotes may also be observed when recombination creates variation in the level of genetic load, for example among offspring after selfing of a single founder (17). Taken together, the propagule model predicts that founder events, genetic drift, inbreeding, gene flow, and natural selection will jointly drive the evolution of SFSs in a metapopulation. However, how SFSs of natural subpopulations evolve in the light of these evolutionary processes is not clear, in particular because these processes are influenced by ecology-related chance events (e.g., immigration and bottleneck size), generating distinct evolutionary trajectories for individual subpopulations (18). At any given time, genetic bottlenecks, hybrid vigor, and extinction events can occur in some, but not other, subpopulations. Despite these local differences, the overall trend of increasing genetic diversity and flattening of the SFS distribution with increasing age of a subpopulation is expected to be shared by all subpopulations but may only be visible in a large enough sample. To investigate the entire range of evolutionary relevant processes, thus, one should study many subpopulations of a metapopulation in parallel and over an extended time period.

The genetic consequences of metapopulation dynamics might be most obvious in small, unstable habitat patches where local extinctions and (re)colonization occur frequently. Here, we studied 118 occupied habitat patches of a dynamic metapopulation of the pond-dwelling crustacean *Daphnia magna* on the Skerry Islands of Southwestern Finland, where subpopulations of the cyclical parthenogen occur in distinct, small water bodies (Figure 1). In this metapopulation, *D. magna* undergoes about six to ten asexual generations annually (planktonic or active phase) and one or two sexual generations annually. Sexual reproduction results in the production of resting stages, allowing *D. magna* to survive the inhospitable winters. An ongoing study with biannual censuses (19–22) has identified high extinction–(re)colonization dynamics in this metapopulation, with about 20 % of all ponds in the study area containing *D. magna* subpopulations, about one fifth of which going extinct every year. This pattern is balanced by a yearly colonization rate of about 5 % for empty ponds. Colonization involves one or a few individuals, most likely originating from a single source population (23). Thus, subpopulation founding is common and associated with severe genomic bottlenecks that might lead to low genomic diversity in newly founded subpopulations compared to older subpopulations with increased genomic diversity (23). Colonization under the propagule model assumes that stochastic processes like genetic drift drive the evolutionary process of the metapopulation (24), although deterministic processes like selection for heterozygotes (through hybrid vigor (16) and because of variation in the level of genetic load (25)) and local adaptation (26–28) are also expected to shape allele frequency dynamics. The dynamic nature and well-studied ecology of the *D. magna* metapopulation make it a particularly suitable model to investigate the dynamics of genome-wide allele frequency changes in dynamic metapopulations.

**Figure 1:**
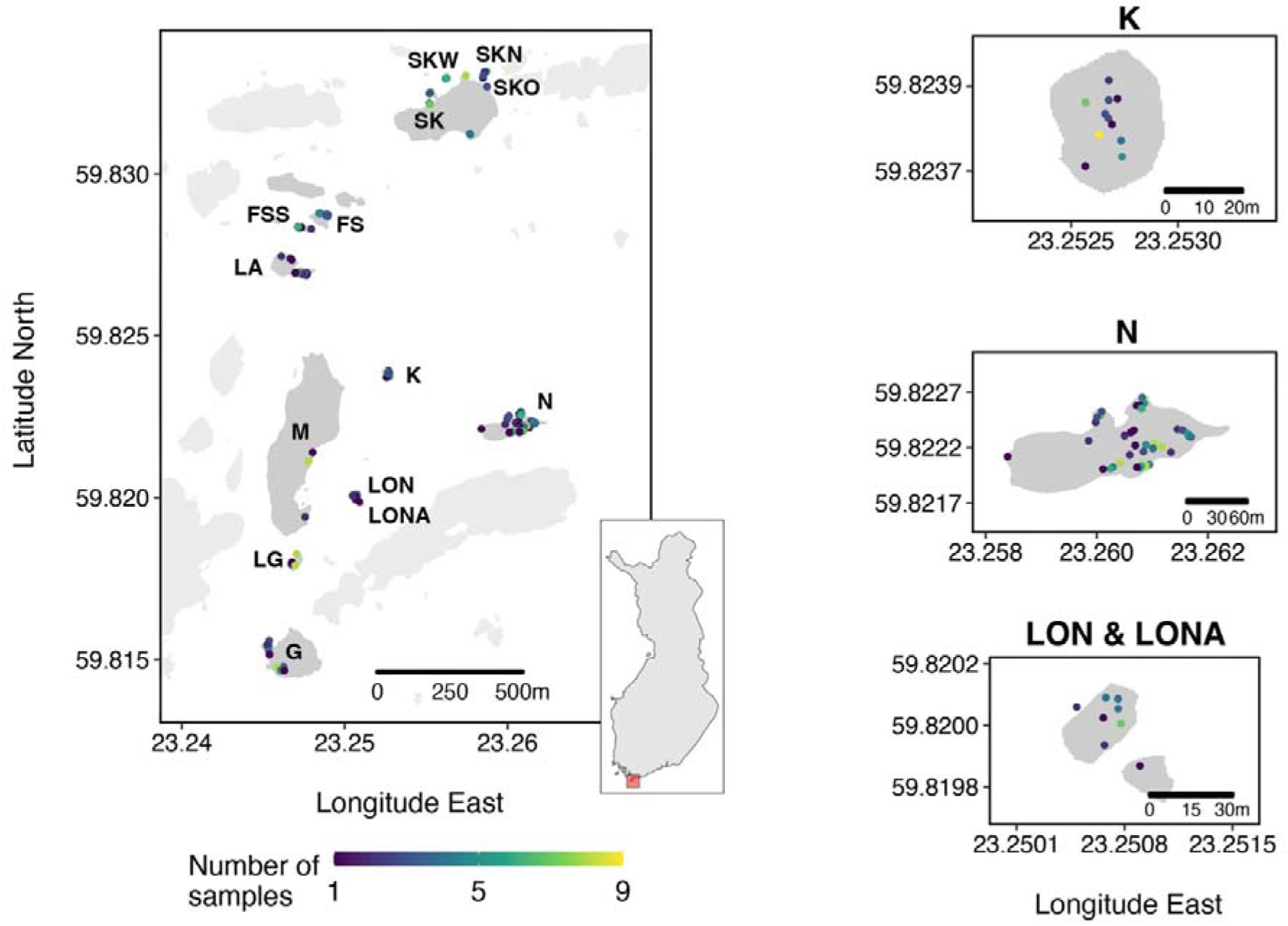
Geographic locations of sampled ponds. The map on the left gives an overview of the study area. The inset map in the middle shows Finland with the study area indicated in red. Islands K, N, and LON/LONA are enlarged as insets on the right to better illustrate the spatial arrangement of individual ponds. Island names and abbreviations were adopted from the long-term study (see e.g. (24)). Ponds are colored according to the number of samples taken during the five-year study period. The study area contains about five times more habitat patches (ponds), but *D. magna* did not occur there during the study period. Islands colored light gray are not part of the biennial census and were, therefore, not sampled.

This population genomic study aimed to investigate the evolutionary process of a dynamic metapopulation by biannually sampling subpopulations of different ages and ecology for five consecutive years. Previous studies on this metapopulation were based on single cross-sections and did not follow the local dynamic across time (23,24,26). Instead, temporal dynamics were studied indirectly by regressing patterns against the recorded population age. This approach is limited, however, as older populations occur in more stable habitat patches and are, thus, a biased subsample of all subpopulations (19). Our new approach—to quantify the distributions of SFSs in individual subpopulations and follow them over time—disentangles the roles of the distinct evolutionary forces that drive genomic changes in dynamic metapopulations. With a five-year sampling period of all subpopulations in a metapopulation with over 550 habitat patches, we are able to witness subpopulation founding and gene flow events with subsequent hybrid vigor in subpopulations of diverse ages. To supplement the analysis of our natural samples, we also conducted simulations of the population genetic processes across time. Together with previous findings from empirical and experimental studies on the same metapopulation, these approaches seek to understand the effects of founder events, natural selection, dispersal, hybrid vigor, and the impact of ecological factors on metapopulation evolution. While these scenarios have been described previously as separate events, here we quantify their frequency and individual contribution to the overall dynamics in order to answer questions about how metapopulations evolve and how this differs from expectations for classical populations.

## Results

### Samples and sequencing

We sequenced 418 pooled *D. magna* samples from 118 ponds (Figure 1, mean per pond: 4) across 14 islands (range per island: 1 to 142 samples, mean: 30) biannually from 2014 to 2018 (range per year: 53 to 108, mean: 84). One hundred ninety-six of these samples were collected in spring and 222 in summer. Sequencing resulted in average whole-genome coverages for the *D. magna* reference genome mostly well above 20× (Table S1). After variant filtering, 3,084,861 single nucleotide polymorphisms (SNPs) were retained for the analysis.

In individual subpopulations, alleles and their frequencies are not expected to change quickly unless specific events like hybrid vigor after gene flow are present. It has been suggested that hybrid vigor is important in shaping SFSs in focal metapopulations, and its effects on allele frequencies in subpopulations—namely a high pairwise *F*_ST_ and an increase in the number of heterozygous sites between two consecutive sampling time points—are well understood. By comparing all pairs of consecutive samples, we found 19 subpopulations that matched indications of gene flow plus hybrid vigor (Figure 2, Table 1A). Therefore, in addition to the overall data, we analyzed these subpopulations separately from those presumably without hybrid vigor events. Further, we observed ten extinction–recolonization events in the field, where a subpopulation went extinct and a new subpopulation recolonized the same pond. Subpopulations of the pond before and after these extinction– recolonization events were treated as separate subpopulations. We calculated pairwise *F*_ST_ between them to detect potential false positives (little change in allele frequencies unnoticeable in the SFS distribution; n = 3; Table S2) and to visualize true positives (strong change in allele frequencies affecting the shape of the SFS; n = 7; Figure 2, Table 1B and S2). Thus, we had a total of 125 subpopulations in our dataset.

**Figure 2:**
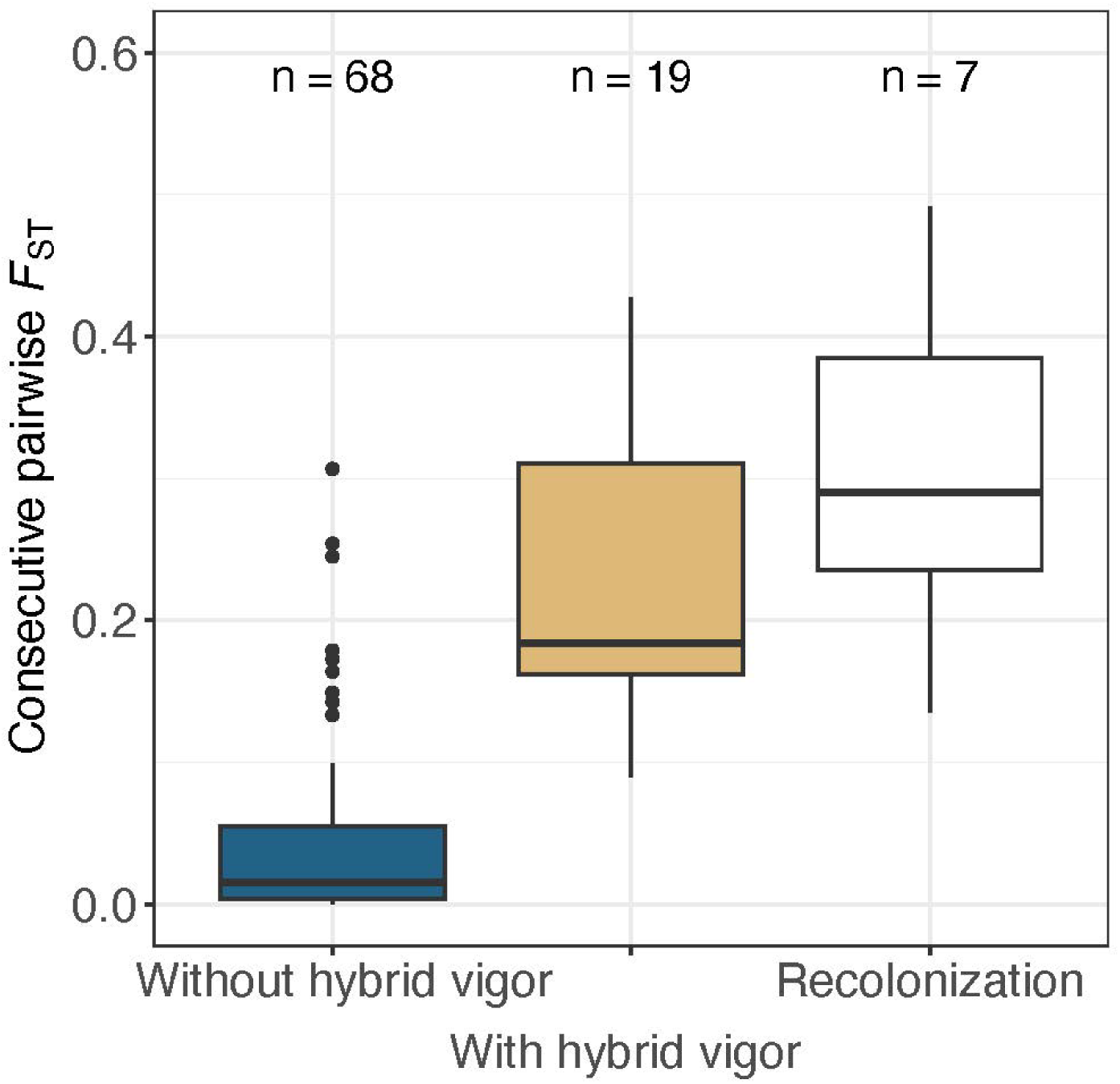
Classification of subpopulations into groups undergoing different processes. Subpopulations with high *F*_ST_ between two consecutive samples and a simultaneous increase in heterozygous sites were classified as having gene flow plus hybrid vigor. The figure shows pairwise *F*_ST_ for consecutive subpopulation samples with hybrid vigor events (yellow) and pairwise *F*_ST_ for consecutive samples without hybrid vigor (blue). The latter is generally low but includes a few high *F*_ST_ values for subpopulations where we observed no increase in heterozygous sites. High *F*_ST_ without a gain in heterozygous sites might stem from population bottlenecks or from extinction–recolonization that happened too quickly to be detected by our sampling (cryptic extinction–recolonization). The right box (white) shows pairwise *F*_ST_ between subpopulations of the same pond before and after a recorded true positive extinction–recolonization event. This and subsequent analyses focused on the temporal dynamics only use subpopulations with at least two years of sampling success.

**Table 1:**
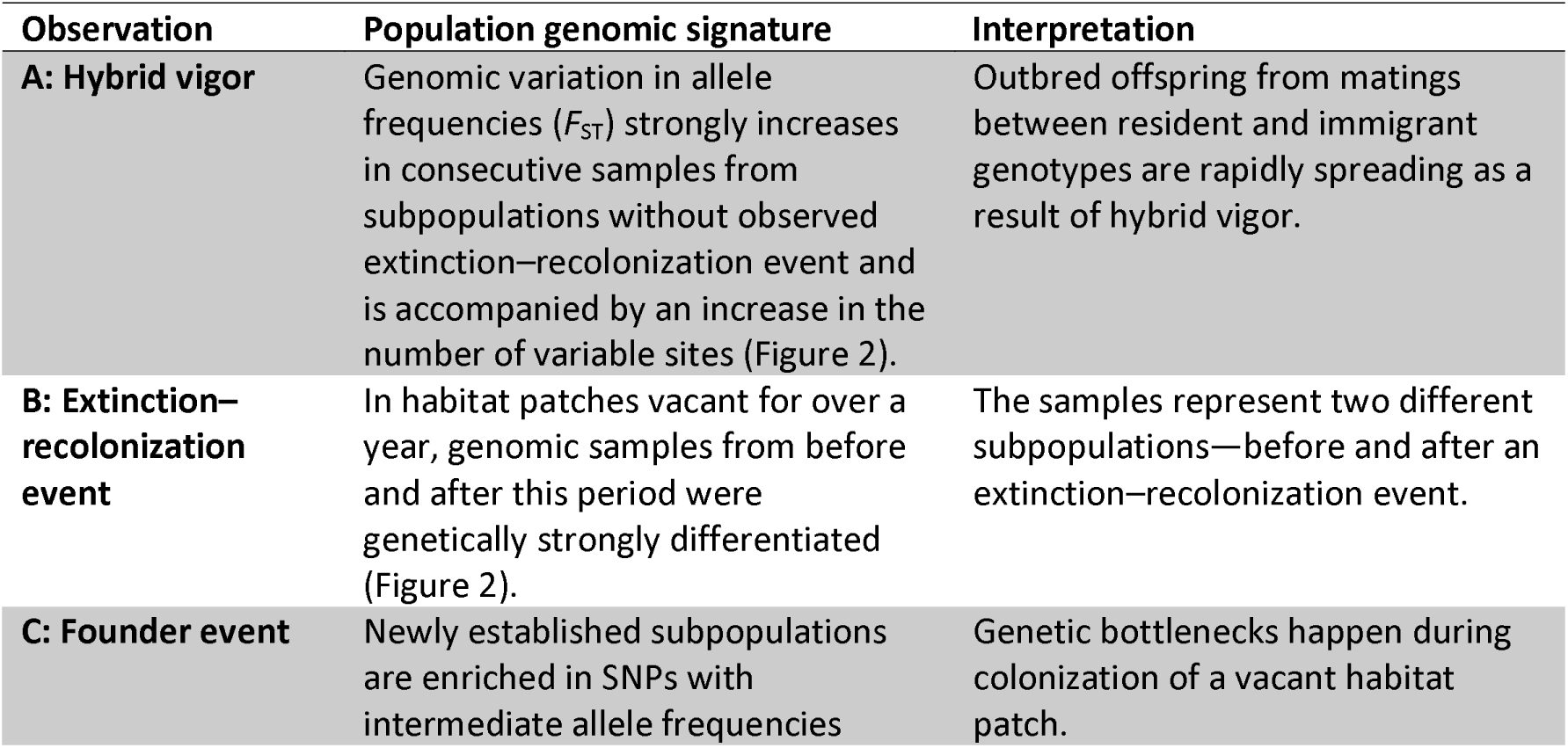
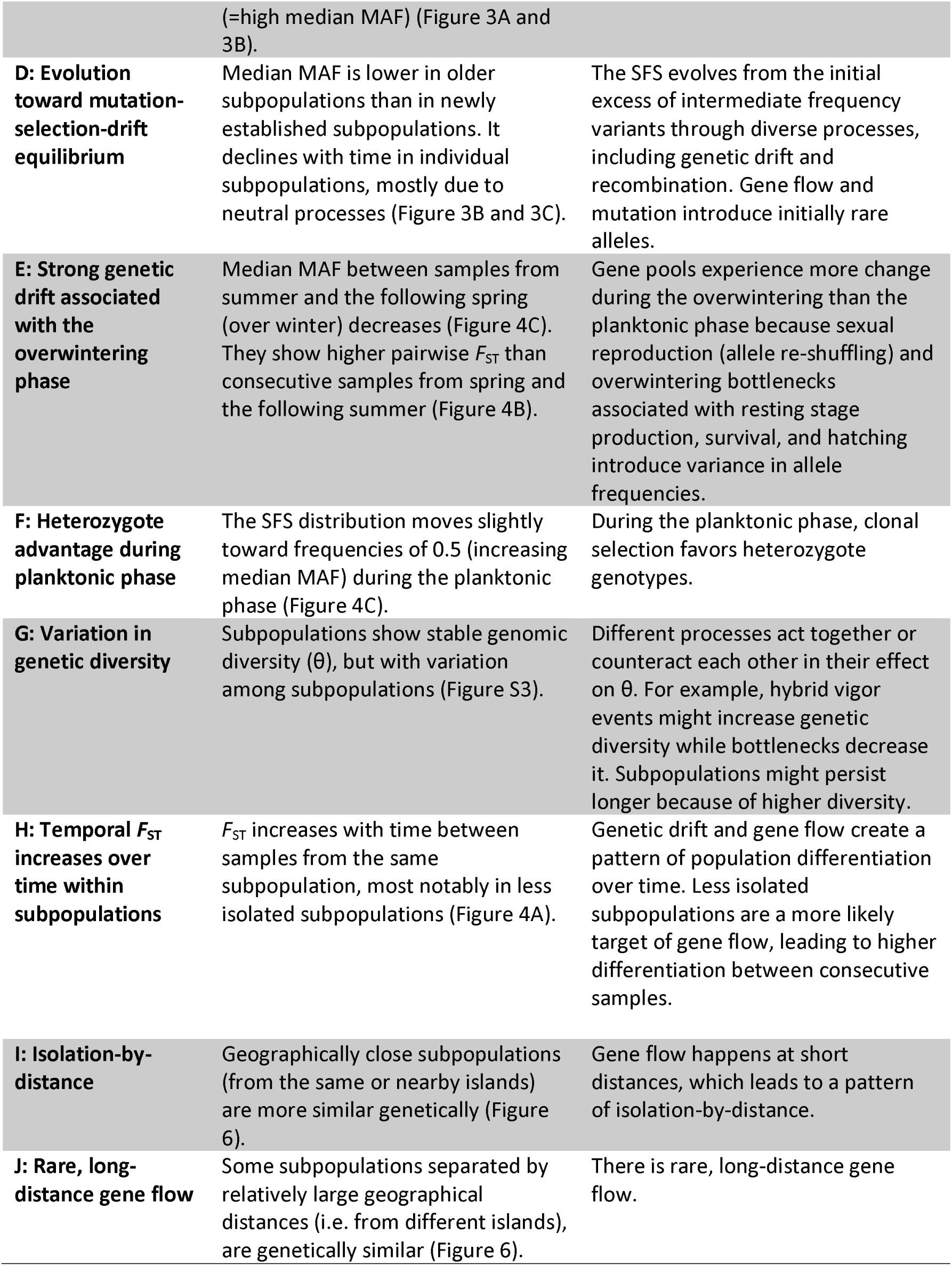
Summary of biological dynamics inferred through the analysis of genomic summary statistics in the metapopulation.

### Shifts in genome-wide allele frequencies

We expect the folded SFS distribution of a subpopulation to generally evolve from left to right skewed with increasing age. However, this prediction is based on neutral evolutionary processes. Deterministic processes may push the SFS distribution (back) in the opposite direction to become more left skewed. As a simple summary statistic for describing the distribution of the SFS we use the median frequency of minor alleles (median MAF), because it is relatively unsensitive to the exact shape of a SFS. As a first approach to investigate whole-genome allele frequency distributions in subpopulations, we tested whether median MAF was associated with the age of the subpopulation by comparing subpopulations of different ages. We found that SFSs of young subpopulations showed higher median MAFs (more SNPs with intermediate frequencies, i.e. left skewed) than older subpopulations (see SFSs in Figure 3A, Table 1C and 1D, statistics in Figures 3B and S1; young versus intermediate: *t*(87.131) = 3.5602, *p* = 6.029 × 10^-4^, young versus old: *t*(76.424) = 5.8076, *p* = 1.372 × 10^-7^, intermediate versus old: *t*(63.114) = 1.3772, *p* = 0.173; mixed-effect model: *F* = 26.73, *p* = 6.16 × 10^-7^).

**Figure 3:**
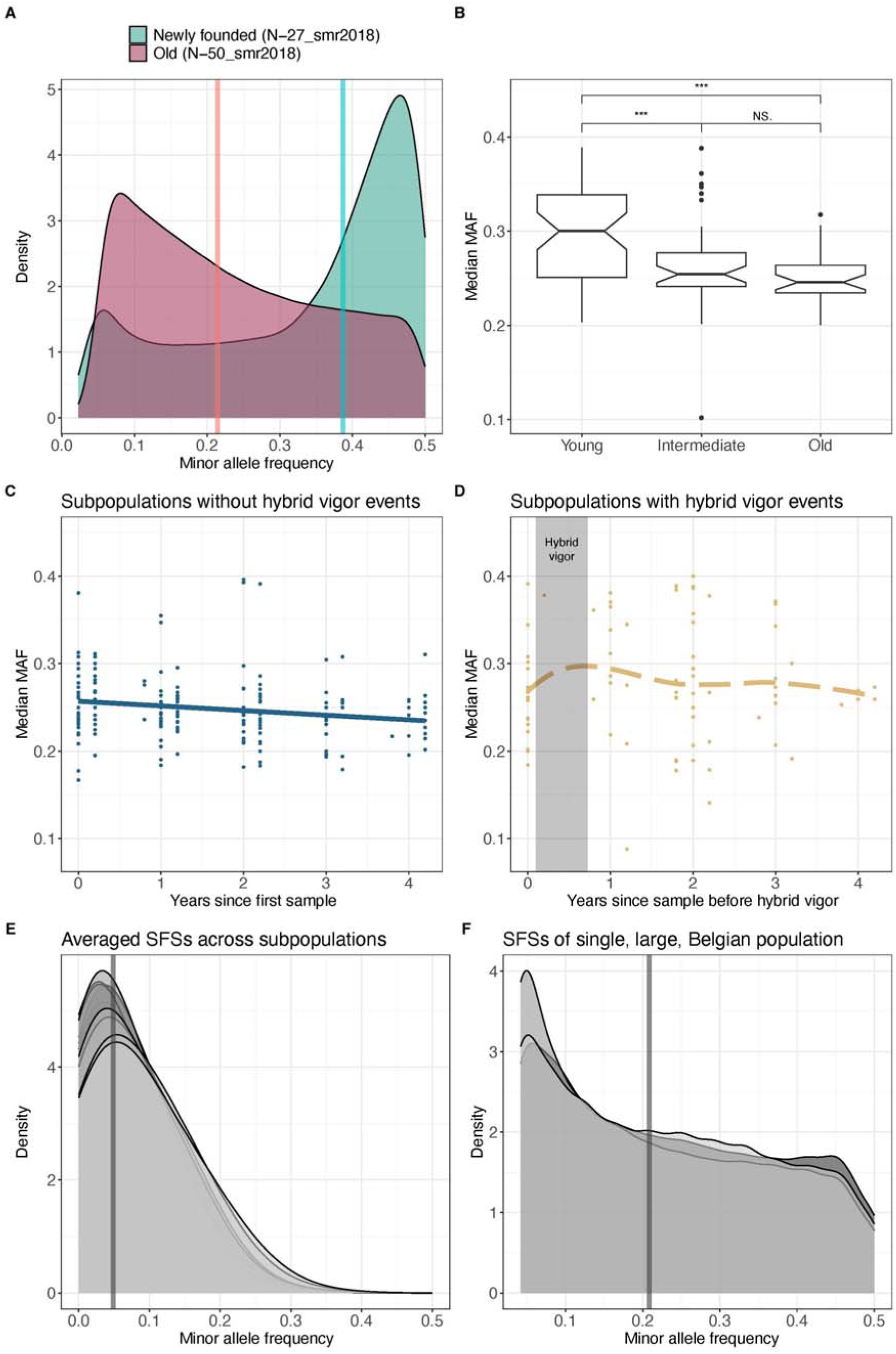
Minor allele frequency distributions and their median in subpopulations of different age and status. (A) Example of a newly founded subpopulation with a left skewed SFS distribution vs. one of the oldest subpopulations (age = 36 years) with a right skewed SFS distribution. (B) On average, median MAF is higher in younger subpopulations than in older ones. Age class definitions are from Angst et al. (24): Young: ≤ 2 years (≃ newly founded, n = 50), Intermediate: > 2 and ≤ 15 (n = 41), and Old: > 15 (= most stable, n = 33). (C) Median MAF decreases in individual subpopulations during the study period. The regression line is from a simple linear model. (D) Subpopulations that showed a hybrid vigor event (indicated with a shaded area) deviate from those without such an event by an initial increase in allele frequencies, as revealed by local regressions with LOESS (dashed line). (E) In the overall metapopulation, the median MAF is stable, and the SFS distribution resembles a classical population. Each of the ten curves represents allele frequencies averaged across all subpopulations at a given sampling point. Alleles close to a frequency of 0.5 are rare, indicating that only a few alleles are common in all subpopulations. (F) In a single, large, stable population from Oud Heverlee, Belgium median MAF is stable and similar to that of old subpopulations from the metapopulation. Each of the three curves is from a different population sample (n = 12), separated by about nine years. The smallest possible allele frequency for these samples is 1/(2×12). In A), E), and F), bold vertical lines represent the median MAF. Colors in C) and D) are used as in Figure 2.

In following the genome-wide allele frequencies of individual subpopulations for up to five years, we found an overall decrease in median MAF over time, i.e., the SNPs’ minor allele frequencies decreased (*F* = 4.54, *p* = 0.034; Table 1D). This decrease was mostly due to allele frequency changes over winter (in the resting stage phase), not during the planktonic phase in summer, where we observed a slight increase in median MAF (Paired Wilcoxon: *V* = 570, *p* = 0.011; Figures 4C and S2, Table 1E and 1F). The decrease of median MAF was stronger in the subset of young subpopulations (*F* = 6.89, *p* = 0.011) and in subpopulations without hybrid vigor (*F* = 5.70, *p* = 0.018; Figure 3C) than overall. This effect was not observed in subpopulations with a genomic signature of gene flow plus hybrid vigor, where we expected and observed an increase in median MAF (Figure 3D, Table 1A).

**Figure 4:**
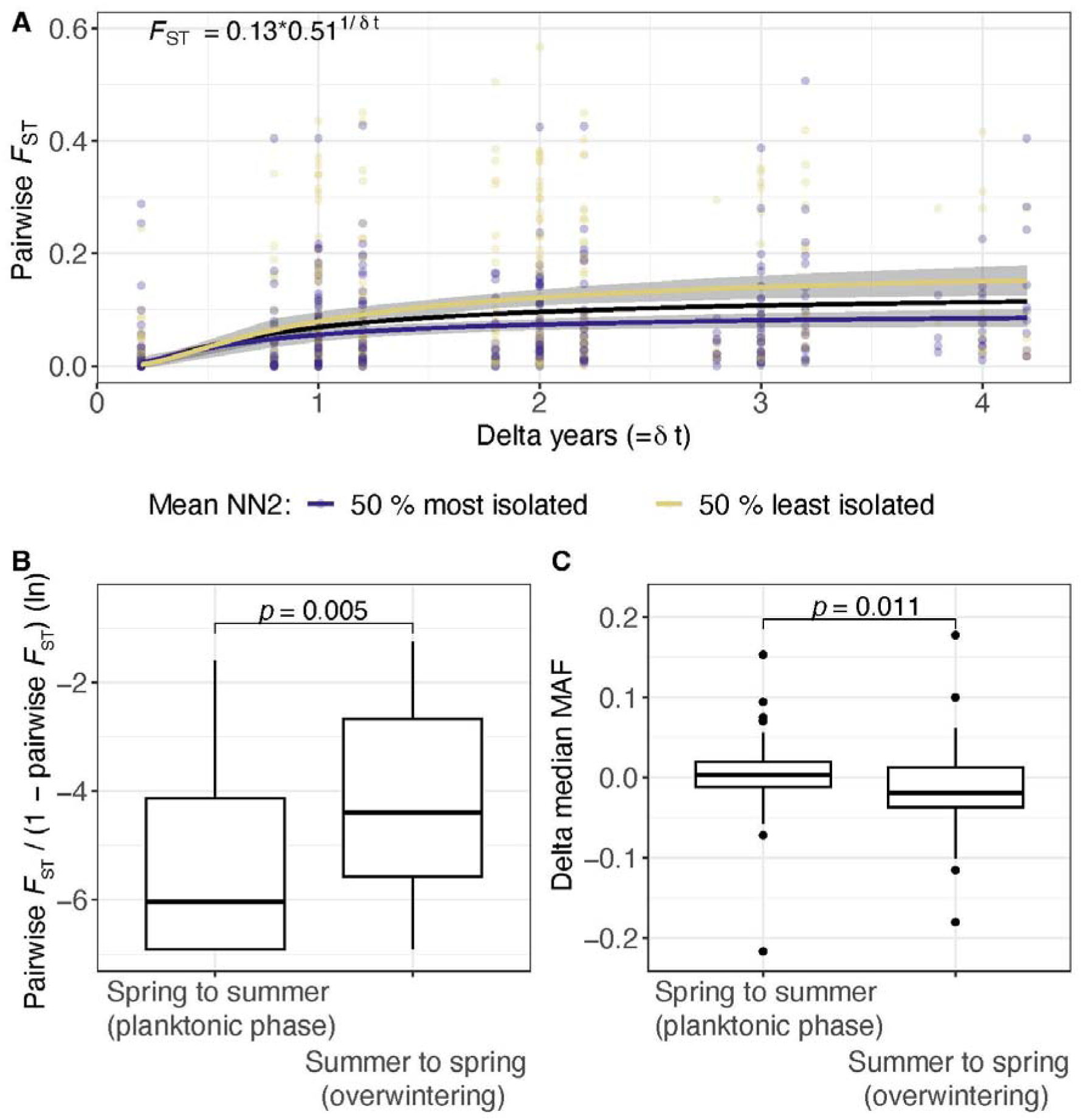
Genome-wide differentiation among temporally separated samples measured as pairwise F_ST_. (A) Overall data and two categorial subsets of subpopulations based on degree of isolation measured as the mean distance to the nearest neighboring subpopulations (NN2). Lines represent predicted *F* values based on nonlinear regression (*F* = ab^1/∂t^) as used in Bergland et al. (32) with 95 % confidence intervals in grey. Statistics describe the model from the overall data (black). The line for the more isolated ponds (50 % most isolated) is lower than for the less isolated ponds (50 % least isolated), indicating less pronounced changes in observed allele frequencies, possibly due to lower local gene flow. (B) Pairwise *F*_ST_ between summer and the following spring samples is larger than between samples from the same year. Untransformed median pairwise FSTs of the two groups are 0.012 and 0.001, respectively. (C) Median MAF decreases on average between the summer and the following spring samples (delta median MAF = -0.013), unlike between samples from the same year where median MAF increases slightly (delta median MAF = 0.005). In B and C, individual groups were compared with paired Wilcoxon tests.

Even though whole-genome allele frequency distributions of individual subpopulations diverge from those expected in a classical population (some strongly), the overall metapopulation might diverge less. This pattern arises because–analogous to individuals in a classical population–individual subpopulations likely have private alleles and only a few alleles are shared among most subpopulations. To examine this, we estimated the allele frequencies on the metapopulation level. We averaged the allele frequencies of all subpopulations, obtaining a distribution similar to what would be expected for a classical population (Figure 3E). To compare the evolution of our composite subpopulation’s genome-wide allele frequency distribution with a single large population of the same species, we used temporal data (3 samples across about 18 years) from the Oud Heverlee *D. magna* population in Belgium (29). Based on the analysis of 2,092,135 SNPs, we estimated that the median MAF of this large population has been stable at 0.2 for about 18 years (Figure 3F). This estimate is lower than the median MAF of the old subpopulations of the metapopulation (about 0.25, Figure 3B).

### Genomic diversity and differentiation over time

Genomic diversity (θ) of individual subpopulations did not generally increase over time (Figure S3). The only subpopulations that showed an increase were those with gene flow plus hybrid vigor, which was part of how we detected this subset (Table 1G). When modeling θ, the interactive effect between the explanatory factors time and isolation was weak (NN2) (*F* = 3.39, *p* = 0.067), suggesting that temporal changes in θ could only marginally be caused by the subpopulation’s degree of isolation. In the pairwise comparisons of consecutive samples, the difference in θ between spring samples and those from the following summer was indistinguishable from the difference in θ between summer and next year’s spring samples (Figure S4).

Population differentiation, estimated as pairwise *F*_ST_ between samples of the same subpopulation, increased with time between samples, ∂t, overall, and in both isolated and less isolated samples (Figure 4A, Table 1H). A nonlinear model described by *F*_ST_ = ab^1/∂t^ fit the observed data better than a linear model, i.e., *F*_ST_ = a + b∂t, (AIC non-linear: -1,436.778, AIC linear: -1,422.533). Samples of the same pond were less differentiated if taken from isolated ponds than taken from less isolated ponds, likely because a main source of variation, gene flow, is higher in the less isolated ponds (Figure 4A). Pairwise *F*_ST_ estimates between consecutive samples of the same subpopulation were three times higher between summer and the following year’s spring samples, i.e., across the winter (on average about 0.012), than between spring and the following summer samples (0.001; Paired Wilcoxon: *V* = 191, *p*= 0.005; Figures 4B and S2, Table 1E), indicating stronger evolutionary dynamics across the winter than the summer.

Regarding spatial population differentiation, pairwise *F*_ST_ between all 418 samples revealed clusters of low differentiation encompassing samples from the same subpopulation and island (Figure S5). This result coincides with previously described patterns of isolation-by-distance (IBD) (24,30,31), indicating that geographically proximate samples are typically less differentiated than more distant samples (Table 1I).

### Simulations

Our simulations focused on isolating the effects of individual processes to help understanding how they shape the SFS. We simulated newly founded populations, tracking changes in their allele frequency under different scenarios for 36 years, the age of the oldest subpopulations in our empirical dataset. For median MAF, we observed the same patterns as in natural subpopulations: in the base simulation, i.e., without immigration or bottlenecks, median MAF decreased with population age (-0.004 per year; Figure 5A). This decrease was stronger in the early years (-0.033 per year), which corresponds to young subpopulations in the empirical data, and during the winter (with sexual reproduction) (mean = -0.013) than during the planktonic phase (only asexual reproduction) (mean = 0.006; Figure 5B). The winter decrease in median MAF can be attributed to increased allele frequency dynamics caused by sexual reproduction and genetic drift. In contrast, the increase in median MAF is caused by clonal expansion (for which there is greater potential after overwintering bottlenecks) and clonal selection. Investigating the effects of genetic drift and newly arising mutations, we found that median MAF in larger populations was initially more stable, close to frequencies of 0.5, suggesting less impact from genetic drift. Subsequently, however, median MAF decreased more in larger populations, because of the presence of more novel mutations, compared to smaller populations (Figure S6).

**Figure 5:**
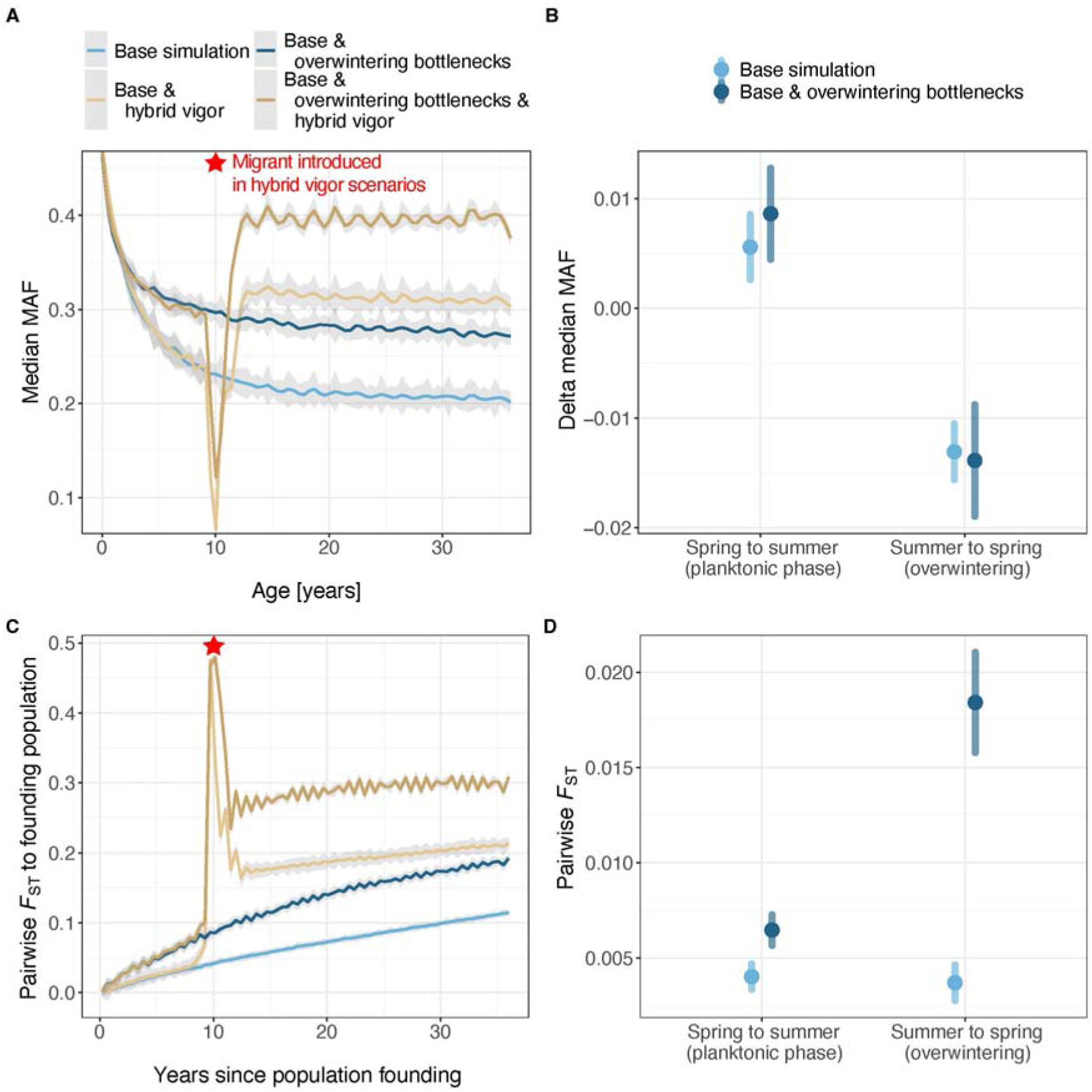
Simulations of processes relevant to the propagule model adapted to the D. magna metapopulation. (A) Median MAF decreases with population age, especially in the early years (base simulation). Higher median MAF values are found if rare alleles are removed by overwintering bottlenecks (base & overwintering bottlenecks) and/or if the alleles of immigrants spread through hybrid vigor (base & hybrid vigor). Migrant(s) are introduced in the hybrid vigor scenarios after ten years, indicated with a red star. (B) The decrease in median MAF is mainly associated with overwintering. In contrast, clonal selection likely increases median MAF during the planktonic phase. (C) The evolution of pairwise *F*_ST_ of a population, relative to its founding population, under the different scenarios outlined in the legend of Figure 5A suggests a temporal change in genome-wide allele frequencies, especially in the hybrid vigor scenario. (D) Populations are more different before and after overwintering bottlenecks (if they occur) than at the beginning and end of the planktonic phase. A) and C) depict cubic spline regression lines along with 95 % confidence intervals; the error bars in B) and D) represent 95 % confidence intervals around the mean.

In the simulated scenario focused on hybrid vigor (with migration), successful immigration was observed in about half of all replicates, leading to an average increase in median MAF of 0.065 within two years after migrant introduction. Upon the immigrant’s arrival, we observed a steep trough in median MAF because even though mutations of the unrelated immigrant are rare in the population, they make up a significant portion of all mutations (Figure 5A). Observing such a trough in a natural subpopulation sample would be unlikely because immigrant alleles are too rare to be picked up by empirical methodology, and immigrants in nature are likely more genetically similar to residents (they mostly arrive from nearby ponds) introducing fewer initially rare alleles than in the simulated scenarios. Immigrant alleles in the empirical samples would likely be detected only after selection for hybrids raised their frequencies. After being rare initially, immigrant alleles increased in frequency and remained at intermediate frequency in our simulations. This is because the population becomes dominated by hybrids, with deleterious mutations from both parents being masked in a recessive state.

Regarding allelic variation, the simulated data supports our empirical finding that there is a nonlinear temporal increase in pairwise *F*_ST_ (AIC non-linear: -387,258.7, AIC linear: - 383,872.5; Figure 5C). In the hybrid vigor scenario, we observed a sharp peak in pairwise *F*_ST_ when comparing the population at the time of the migrant’s introduction to the time of its founding. This pattern corresponded to the concurrently observed trough in median MAF. While pairwise *F*_ST_ in the founding population increased upon immigrant introduction, a decrease then follows because some immigrant alleles are lost, and the resident population’s allele frequencies are pushed back to frequencies close to 0.5 because of selection for the hybrid offspring (hybrid vigor), making them slightly more similar to the founding population. In populations without successful gene flow, pairwise *F*_ST_ decreases because the immigrant alleles are disappearing. Our simulations further suggest that genetic bottlenecks associated with overwintering, e.g., when only a portion of animals hatch from resting stages or when members of the parent population contribute unequally to the resting egg pool, enhance genetic differentiation over winter: simulated pairwise *F*_ST_ between consecutive summer and spring samples (mean = 0.018) was higher than simulated pairwise *F*_ST_ between consecutive spring and summer samples (mean = 0.006) only in a scenario with bottlenecks over winter, not in scenarios without bottlenecks (both means = 0.004; Figure 5D). In the simulated populations, θ was stable except for the hybrid vigor scenario, where we observed an almost three-fold average increase.

### Population structure

As observed earlier (24), the overall structure of our larger genomic dataset reflected the spatial structure of the focal metapopulation: most geographically close samples (from the same pond or island) formed clusters (Figure 6). Also, samples from close neighboring islands, such as LON and LONA, as well as SK, SKN, SKO, and SKW, formed single clusters. This clustering supports the observation of IBD. However, despite strong clustering by island, we see evidence for inter-island dispersal: some samples from different islands cluster together (Table 1J). For example, island K’s cluster includes a subpopulation each from FSS and LG (Figure 6B), a subpopulation from M clusters with samples from LON and LONA (Figure 6C), and the SK island group clusters with a subpopulation from each FSS and LA (Figure 6D). Furthermore, subpopulations form some islands separate into multiple clusters (e.g., FSS, LA, LG, M, N) (Figure 6), whereby the ancestors of the subpopulations of these subclusters presumably originate from distinct colonization events (Figure S7).

**Figure 6:**
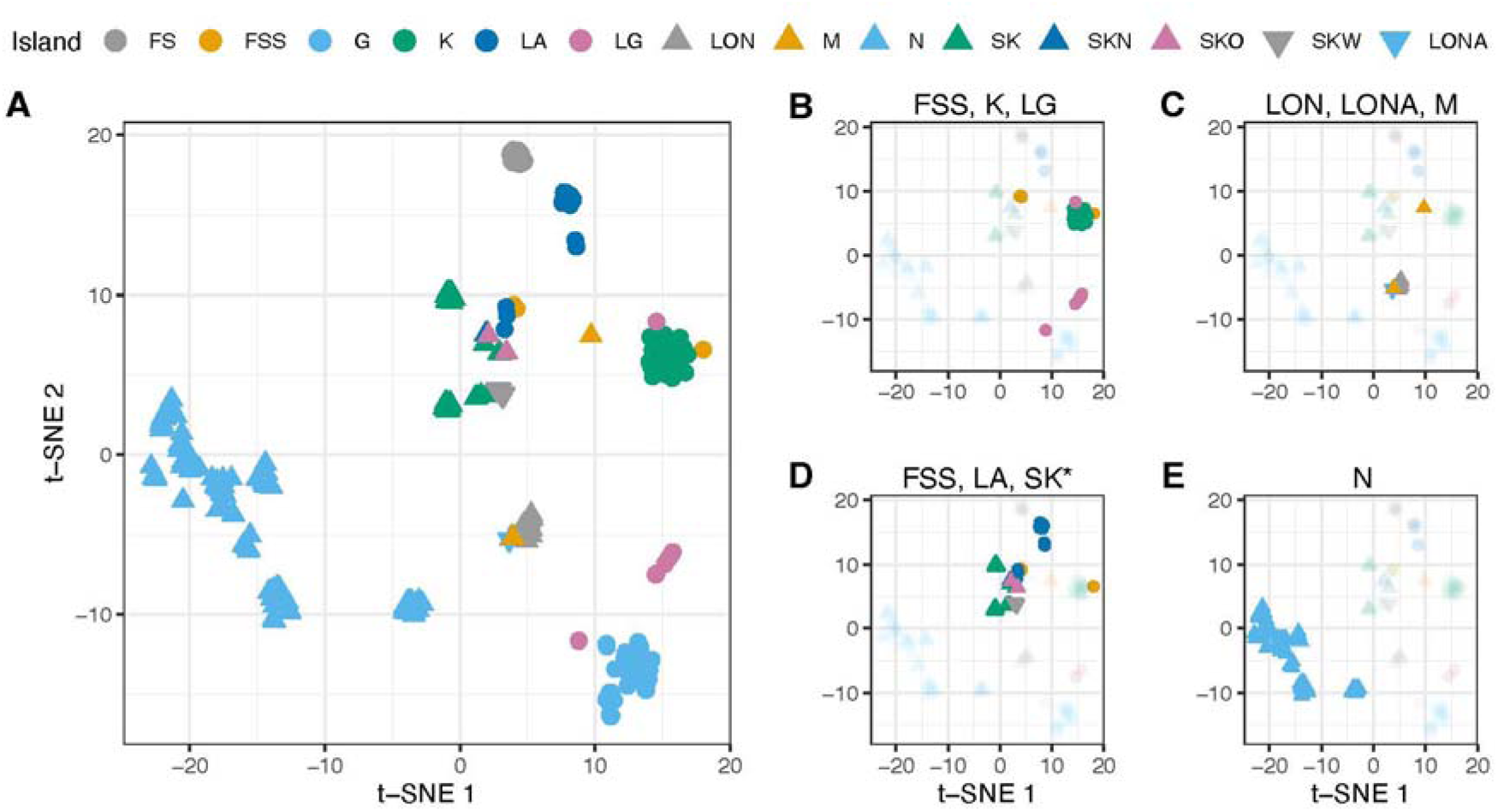
Spatial population structure of the metapopulation. Dimensionality-reduction of whole-genome allele frequencies using t-SNE reveals clustering of samples from the same island in most cases (A). B) to E) highlight samples from different islands clustered together and samples from the same island separated into multiple clusters, indicating long-distance dispersal and distinct colonization events. Shape and color of symbols indicate the island of origin (see legend) and SK* stands for the island group SK, SKN, SKO, and SKW.

## Discussion

We used intensive spatiotemporal genomic sampling to investigate how general and metapopulation-specific processes shape the evolution of genome-wide allele frequency patterns in an ecologically well-studied dynamic metapopulation (19–22). Previous studies have suggested that this metapopulation follows the propagule model (19,20,24,33,34), which this study corroborates. We found that genomic bottlenecks during subpopulation founding skew the distribution of genome-wide allele frequencies to intermediate frequencies. We hypothesized that microevolutionary processes, including gene flow, genetic drift, and newly arising mutations, gradually shift SFS distributions away from intermediate frequencies. Our analysis confirmed this hypothesis and further revealed shifts in the opposite direction in some subpopulations, likely due to the rapid spread of hybrid offspring from inbred resident and immigrant genotypes, i.e., hybrid vigor.

### Deepening our understanding of the propagule model

By unraveling the dynamics of SFS distributions in the *D. magna* metapopulation and by applying numerical simulation, we advance the understanding of how metapopulations evolve and explore the population genomic consequences of the propagule model. Our analyses of both empirical and simulated data quantifies the contribution of previously observed and newly discovered factors in metapopulation evolution and supports the hypothesis of evolution under the propagule model by showing that newly founded subpopulations have high median MAF, consistent with the idea that they are founded by a single or a small number of animals (30). This finding was corroborated by simulations and is also supported by previous studies on turnover dynamics, the number of colonizers, and mitochondrial as well as nuclear genomic data, which have suggested evolution under the propagule model for this metapopulation (19,20,24,33,34). A previously unknown phenomenon is that changes in the local allele frequencies were considerably stronger from summer to the following spring (the overwintering period) than from spring to the following summer. This seasonal pattern highlights the previously unknown role that processes related to sexual reproduction (resulting in resting stages), resting, and hatching play in the evolution of SFSs in subpopulations and, more generally, highlights the advantage of our holistic approach. Sexual reproduction in cyclic parthenogens involves various stochastic aspects, which may lead to unequal contribution of members of the planktonic population to the resting egg pool and to the following planktonic population. For example, genotypes have an unequal propensity to reproduce sexually (35), the production and survival of resting stages can vary, and hatching can be temporally asynchronous. Thus, overwintering may be associated with reductions in the effective population size, genetic bottlenecks, and the previously described increased risks of extinction (19). Furthermore, we found clear evidence for long-distance dispersal. Previously thought to be rare in the focal metapopulation (19,21,22), we observed at least five cases where long-distance dispersal between islands up to 500 m apart is the most plausible explanation (Figure 6). Dispersal also led to hybrid vigor, which we observed in about 22 % of all subpopulations for which multiple samples were available (Figure 2). The observed temporal increase in median MAF in subpopulations with hybrid vigor has been suggested by previous studies (10,16). Our data suggests a hybrid vigor event can occur in a given subpopulation about once every 21 years, or at a chance rate of about 5 % per year. The remarkably similar values of hybrid vigor rate and colonization rate suggest that about 5 % of occupied and unoccupied ponds in this metapopulation are likely to experience successful immigration each year.

Prior to this study, the genetic model of the focal metapopulation was inferred from empirical and experimental studies that provided valuable insights, but also had distinct inferential weaknesses. First, the empirical studies of natural subpopulations were based on single cross-sections, and the effects of time were approximated using subpopulations of different ages (23,24,26). Subpopulation age was thereby confounded with other potentially age-related factors like habitat stability and quality, infection status, and isolation. Second, although experiments demonstrated strong effects such as hybrid vigor, the relative importance of these effects for the overall evolutionary process could not be assessed (10,16). Third, potential confounding factors were controlled for in the experiments and thus did not contribute to the overall variance (16). Fourth, experiments tended to test contrasting conditions, thus potentially overestimating the strength of a factor. For example, Ebert et al. (16) tested the effects of inbreeding and gene flow by contrasting inbred versus outbred individuals and distant immigrants versus residents. Although necessary for the chosen experimental design, this approach did not consider that variation is not bimodal but continuous. Overall, the picture of the evolutionary processes in this metapopulation had been simplified in previous studies, creating the impression of rather strong individual effects. Here, we provide a larger context for those studies—including findings of high turnover dynamics, small numbers of subpopulation founders, low effective population sizes, weak purifying selection, and low rates of adaptive evolution—and present a quantitative picture of metapopulation evolution, as well as compelling evidence for the propagule model, for the strength and frequency of hybrid vigor events, and for the effects of subpopulation aging. Furthermore, by showing the genomic consequences of high extinction–(re)colonization dynamics, this study demonstrates the effectiveness of using genome-wide allele frequencies to study evolution in metapopulations. We conclude that the SFSs of dynamic metapopulations do not match the norms of classical populations, upon which many widely used population genetics models are based.

### Genomic consequences of founder effects and gene flow

In line with the propagule model of population founding, we found SFSs skewed to intermediate frequencies in newly founded subpopulations. The median MAF was higher in younger subpopulations than in older ones, suggesting that the SFS distribution shifts to the left over time, converging to a right skewed SFS distribution. We observed this change in real-time as we followed the genome-wide allele frequencies of subpopulations for up to five years. This is supported by in silico simulations. The use of the median to quantify the distribution of genome-wide minor allele frequencies is robust in that it depends little on the exact shape of the SFS. Given the variety of concomitant biological processes in the focal metapopulation, this simple measure turned out to be informative.

In contrast to the overall pattern of a decline of median MAF over time, this median increased in some of the subpopulations over time, alongside with rapid and strong genomic differentiation and an increase in the number of heterozygous sites. We interpret these observations as gene flow plus hybrid vigor events because such fast changes are strongly suggestive of a deterministic process, particularly if they are visible on a genome-wide scale. Without a selective advantage, alleles introduced by immigration events or mutation would mostly disappear, because of their rareness. Only a few alleles would stochastically rise in frequency over time. Our study could not directly test for hybrid vigor due to the lack of genotype information in our data; however, Ebert et al. (16) speculated hybrid vigor to be common in this metapopulation of *D. magna* and showed hybrid offspring resulting from the mating of residents and immigrants outcompeting parental lineages usually within one season. This strong effect may have been a consequence of their experimental design, however, which had hybrid offspring at a relatively high frequency at the beginning of the season. This scenario is unlikely in natural populations, where hybrid offspring will initially be rare and where, therefore, their spread will take longer. Furthermore, in Ebert et al. (16), the introduced migrants had to differ in marker genes from the residents, so they may have been more differentiated than an average immigrant, which is more likely to originate from a nearby, genetically similar subpopulation. This large differentiation may also have amplified the hybrid vigor effect. Here, we may have missed some hybrid vigor events, as their effects were not visible as clearly. Furthermore, we cannot fully rule out the possibility that we misclassified some subpopulations as having experienced gene flow plus hybrid vigor, that do, in fact, represent false negative (very fast and undetected) extinction– recolonization events. Nonetheless, our method using biannual censuses for identifying extinction–recolonization events has been shown to be robust (19), and the observed substantial increase in the number of heterozygote sites is not expected after a recolonization event.

Cross-sectional samples of this metapopulation, where older subpopulations were more diverse, have suggested increasing genomic diversity (23,24), while our temporal sampling scheme showed no general increase in the genomic diversity of individual subpopulations. Diversity is expected to increase as a consequence of gene flow (plus hybrid vigor), selection for heterozygote genotypes (clonal selection for less inbred genotypes) (25), and de novo mutations. It may decrease as a consequence of genetic drift and population bottlenecks. Selection for heterozygotes is observed here and includes hybrid vigor after gene flow, which was observed only in a few subpopulations, and selection for heterozygotes resulting from variation in local inbreeding. The latter has been used to explain small increases in heterozygosity in rock pool populations during the planktonic phase (25) and is a likely explanation for the here observed increase in median MAF during the summer. The effect of de novo mutations is unlikely visible given the low mutation rates and the short time scale of our study. Genetic drift is a strong force, in particular in small populations and in our *D. magna* metapopulation (24), and may lead to the random fixation of alleles and therefore lower local diversity (36). Genetic drift is even stronger if genetic bottlenecks occur, as seems to be the case during the overwintering of subpopulations. Taken together, the absence of an overall change in genomic diversity over time in our current study reflects the conflicting trends of selection for heterozygotes versus genetic drift and bottlenecks. As these processes differ among habitat patches, they will result in variable evolutionary trajectories across subpopulations. These considerations highlight the difficulties in predicting a general trend for the change of genomic diversity over time. The absence of an increase in genomic diversity over time in the current study, while older subpopulations have been reported to be more diverse than younger subpopulations (23,24) has, however, a different explanation. Older subpopulations are a non-random, small subset of surviving subpopulations. Most subpopulations have a low life expectancy, with about 80 % of subpopulations not becoming older than three years, while some subpopulations survived for as long as the biennial census was done (> 36 years) (D. Ebert, pers. observation) (22). The few old subpopulations are typically associated with larger pond (and thus population) size and higher habitat patch stability, both of which contributes to their higher survival chances (22,37). As a consequence, they may have experienced more hybrid vigor events and less genetic drift and, thus, have higher genetic diversity than a random sample of young subpopulations. Therefore, old subpopulations are a nonrandom sample of all subpopulations, so conclusions drawn from them regarding the evolutionary process are biased.

Our longitudinal sampling revealed that pairwise *F*_ST_ increased with time between samples of the same subpopulation (Figure 4A), though samples from the same subpopulation and island remain more similar to each other than to others (Figures 6, S5, and S7). Bergland et al. (32) likewise found an increase in *F*_ST_ with time between samples in a natural *Drosophila melanogaster* population. They attributed this change to rapid response to selection driven by seasonal changes. For our *D. magna* metapopulation, genetic drift likely contributes to this increase because of its dominant role in this metapopulation and because adaptation to the local environment is thought to be weak (24,26,28). Gene flow also likely contributes to the gradual rise in population differentiation over time between samples from the same subpopulation, as suggested by the steeper rise in less isolated samples with more potential for gene flow (Figure 4A). Our analysis of *F*_ST_ further shows that changes in allele frequencies are more profound after hatching from sexually produced resting stages in spring than after a period of repeated asexual (clonal) reproduction. After hatching, fewer alleles are at intermediate frequencies. In contrast, intermediate frequency alleles are slightly enriched during the planktonic phase, potentially due to clonal selection associated with heterozygote advantage (25). Taken together, these results suggest that the period of sexual reproduction, overwintering, and hatching plays an important role in the evolution of metapopulation SFSs. Seasonal differences have recently been highlighted as key drivers of metapopulation dynamics in a demographic study of the same metapopulation (19).

Our results add knowledge to the *D. magna* system and are generalizable to many aquatic organisms in patchy habitats with a similar life cycle, such as ostracods, copepods, rotifers, and other Cladocera (38–42), many of which also occur in the focal rock pool habitat. We expect many of these systems also evolve under the propagule module and show similar genomic patterns. We further think our findings can be, in part, generalized to many other species with metapopulation ecologies driven by extinction–(re)colonization dynamics, as other major model systems in metapopulation research, for example, the Glanville fritillary butterfly *Melitaea cinxia* and the herb *Silene dioica*, partially follow the propagule model and show patterns of genetic variation consistent with our findings (43,44). Even species without a natural metapopulation structure are affected by extinction and (re)colonization dynamics due to the ongoing extinction crisis (45). This study of the highly dynamic *D. magna* metapopulation thus adds empirical data to the scarce genomic resources of dynamic metapopulations, empirical knowledge to the study of how they evolve, and hopefully the kinds of insights that will aid in their conservation.

### Potential limitations due to metadata and sampling procedure

A potential bias in our analysis might arise from our method for estimating subpopulation age. If we found no animals on three or more consecutive habitat patch inspections, we assumed subpopulation extinction. If we only had one or two visits with no animals, we did not assume extinction to account for the potential presence of undetected animals or of animals present only as resting stages. Although this method has been shown to work well for the focal metapopulation overall (19,21,24,37), some subpopulations may remain undetected for longer periods and, thus, were classified as extinct even though they were not. We tried to correct for such false positive extinction–recolonization events by comparing summary statistics from the genomic data before and after potential recolonization. In so doing, we confirmed seven true, but detected three false extinction events. Still, even after this correction, some young subpopulations show low median MAF, which seems to contradict the propagule model. These could result from subpopulation founding by multiple individuals, either simultaneously or in short sequence (potentially from multiple source subpopulations or the migrant pool model (9)). Sporadic founding by multiple individuals from the same source subpopulation has previously been suggested based on field observations (34).

The other form of age-estimation error occurs when true extinction and recolonization events happen within less than a year. In such cases, we would have estimated subpopulations to be older than they were, which would explain why some intermediate-aged subpopulations show an enrichment of intermediate frequency alleles. Another explanation, apart from hybrid vigor, for intermediate-aged subpopulations with an enrichment of intermediate frequency alleles is the occurrence of strong population bottlenecks. For example, when only very few animals hatch after the resting phase, or when one hatchling hatches much earlier than others and has a head start in asexual propagation. Both, strong genetic bottlenecks and false negative extinction–recolonization events, likely result in high pairwise *F*_ST_ between consecutive samples and a decrease in the number of heterozygous sites, which we observed in about 8 % of the subpopulations (Figure 2).

Further difficulties arose from the field sampling itself. We were not always able to collect the target number of 50 individuals per sample. Some ponds contained low numbers of individuals at the time of sampling, so we had to exclude extant but insufficiently large populations from sampling to avoid disturbing the natural dynamics. Therefore, and because of the extinction–(re)colonization dynamics, some ponds’ time series were not complete. To avoid an imbalanced study design, we focused analyses on temporal dynamics only on subpopulations with at least two years of sampling success.

## Conclusions

Because evolutionary processes in classical populations are, unlike metapopulations, well studied and understood, the assumptions about the population genetics of natural populations are based largely on these stable populations. However, our study highlights that the evolutionary process in dynamic metapopulations with unstable subpopulations deviates strongly from classical populations. This difference is mainly evident in three factors. First, local extinction limits the lifespan of subpopulations and, thus, the degree to which subpopulations can approximate a classical population. Second, (re)colonization of empty patches leads to population bottlenecks and subpopulations with excess alleles at intermediate frequencies, including deleterious alleles. Third, because of genetic bottlenecks, subpopulations attain high levels of homozygosity and genetic load through inbreeding, opening the door for hybrid vigor events after gene flow and for the masking and subsequent purging of deleterious mutations. Together, these factors reduce the efficiency of natural selection, while genetic drift plays a major role in the overall evolutionary dynamics. Metapopulation theory provides a framework–the propagule model–for accurately understanding how these processes affect genome-wide allele frequency distributions (5,9,15). Here, we corroborate the theoretical hypotheses of the propagule model and show empirically how subpopulations in a large metapopulation deviate strongly from our understanding of classical populations. We reaffirm concerns that assumptions like panmixia and Hardy–Weinberg equilibrium for natural populations should be tested before applying common population genetic models to study their evolution (46–48). Dynamic metapopulations evolve differently.

## Methods

### Study system

In this study, we used samples from natural *Daphnia magna* populations inhabiting shallow rock pools (depressions in the bare rock) on the Tvaerminne archipelago in Southwestern Finland (59°50II N, 23°15II E). These rock pools have a mean volume of about 200-300 L and will be referred to as ponds in this paper, so as not to confuse them with the term “pool” from a genomic pool-seq sample. *Daphnia magna* is a planktonic freshwater crustacean whose mentioned populations form a metapopulation with extinction–(re)colonization dynamics and gene flow (19,20,22). More than 550 ponds, most of which freeze solid in the winter and many of which dry up during summer, have been surveyed biannually since 1982 for the presence of *D. magna* (20,22). Sexual reproduction in the cyclical parthenogenetic *D. magna* results in resting eggs that enable local populations to survive the inhospitable winter freezes and summer droughts. Furthermore, the resting eggs, which are dispersed by wind, water, and birds, are crucial for migration and colonization of vacant habitat patches.

### Samples and sequencing

We collected *D. magna* from all occupied ponds in the core sampling area of the long-term study area during regular biannual visits in late May/early June (= spring samples) and late July/August (= summer samples) from 2014 to 2018 (Figure 1, Table S1). Collecting individuals, preparing samples, and extracting DNA followed the procedures described in Angst et al. (24). Briefly, we randomly sampled 50 individuals per pond for whole-genomic sequencing of pooled individuals (pool-seq). We reduced non-target DNA (gut content and microbiota) by applying an antibiotic treatment and inducing gut evacuation. DNA integrity during transport was preserved by adding RNAlater (Ambion); for DNA extraction, we used the Qiagen GenePure DNA Isolation Kit supplementing glycogen (Sigma-Aldrich). Library preparation and sequencing were done by the Quantitative Genomics Facility service platform Basel (D-BSSE, ETH), Switzerland using Kapa PCR-free kits and an Illumina HiSeq 2500 and Illumina Novaseq 6000 sequencer, respectively. Genomic samples from the spring 2014 (NCBI database; Bioproject ID: PRJNA862292) were used in a previous study (24).

### Mapping genomic reads and variant calling

We assessed the quality of the raw paired-end sequencing reads using FastQC v.0.11.8 (http://www.bioinformatics.babraham.ac.uk/projects/fastqc) and trimmed them to remove low-quality sequences and adapter contamination using Trimmomatic v.0.39 (49). All bioinformatical software were used with default settings unless stated otherwise. We ran FastQC again to confirm successful trimming and interleaved trimmed reads using seqtk v.1.2 mergepe (https://github.com/lh3/seqtk). We used a snakemake (50) workflow described in Angst et al. (24) (https://github.com/pascalangst/Angst_etal_2022_MBE) to process interleaved reads to a VCF file and used the XINB3 individual genome (version 3.0; BioProject ID: PRJNA624896; Fields et al., in prep.) as the reference genome. This genome originates from a genotype collected from our study area and is therefore closely related to the samples used in our study. The snakemake workflow uses the following software: bwa-mem2 v.2.2.1 (51) for mapping reads, SAMtools v.1.7 (52) for file manipulations, Picard Toolkit v.2.23.9 (53) for adding read groups and identifying duplicates, and GATK v.3.8 (54,55) for realigning INDELs and calling SNPs using the UnifiedGenotyper. Furthermore, we estimated the average read depth of each sample using SAMtools function depth. These calculations were performed at sciCORE (http://scicore.unibas.ch/) scientific computing center at the University of Basel. We filtered the resulting VCF file to include high quality sites (QUAL > 30, MQ > 40, QD > 2.0, FS < 60) with biallelic SNP variants (i.e., excluding INDEL variants) using vcffilter from the C++ library vcflib v.1.0.0_rc2 (56) and VCFtools v.0.1.16 (57). Because GATK includes uninformative reads in the depth estimate but not in the allelic depth estimates, we recalculated depth estimates based on the allelic depths using VcfFilterJdk v.1f97a34 (58). Afterward, we masked entries with DP less than ten and larger than twice the mode, calculated for each sample separately using VCFtools and BCFtools v.1.9 (59) and masked such with AD of the minor allele equal to one using VcfFilterJdk. This minor allele read count filter is, in pool-seq studies, a more conservative, coverage-aware filter to avoid sequencing errors than the more commonly used minor allele frequency (MAF) filter (60).

When examining the sequencing data, we noticed that the mapping percentages of genomic sequencing reads for some samples was comparatively low (Table S1). This pattern was not unexpected, given that many samples contained a natural parasite of *D. magna*, the microsporidian parasite *Hamiltosporidium tvaerminnensis* (61). Unfortunately, this resulted in low coverage of the host’s genome in four samples, which we therefore had to exclude from the analysis (i.e., N-71_spr2014, LON-1_smr2014, N-61_smr2014, and N-89_smr2014; Table S1). Two samples failed the sequencing entirely (i.e., N-71_spr2015 and SKW-1_spr2015; Table S1).

### Population genetic analyses

We estimated genomic diversity, θ, by averaging estimates of 100 Kbp windows from grenedalf v.0.2.0-beta (62), a recent reimplementation of PoPoolation (63). We chose θ because its calculation is less affected by the SNPs’ allele frequencies than π. Genome-wide allele frequencies were analyzed separately. To investigate spatiotemporal population differentiation, we estimated pairwise genomic differentiation (pairwise *F*_ST_) between subpopulations using the R package *poolfstat* v.2.0.0 (64) in R v.4.2.2 (65). From *poolfstat*’s pooldata object, we also estimated allele frequencies and retained positions with low missingness for conducting t-SNE using the *Rtsne* v.0.15 (66) R package with nine dimensions kept from the initial PCA step and 5,000 iterations.

During our sampling, we noted ten extinction–recolonization events where a previously occupied pond contained animals after being vacant for at least three consecutive visits. The likelihood of not detecting a subpopulation for three consecutive visits has been estimated at below two percent, as the detection probability of *D. magna* is 0.74 in this survey (19). Seven of these ten recolonizations were true positives based on our interpretation of the genomic data, i.e., they showed high pairwise *F*_ST_ and SNP turnover before versus after recolonization (Table S2). Thus, we treated samples from before and after recolonization events in these seven ponds as separate subpopulations in our analyses. For the three (apparently) false positive recolonization events, the four samples collected after the event showed little change in allele constitution compared to samples from the original subpopulations, so it was not clear if they stem from a separate subpopulation and were excluded from analyses requiring a (relative) age estimate. By following *F*_ST_ and the gain/loss-ratio of heterozygous sites over time, we further classified subpopulations into those with gene flow plus hybrid vigor and those without. Gene flow in combination with hybrid vigor is expected to cause comparatively strong differentiation between two consecutive samples as well as an increase in heterozygous sites. For analyses focusing on understanding temporal changes of whole-genome allele frequencies within subpopulations, we only used subpopulations whose sampling period spanned at least two samples from two years. Subpopulations where we could only obtain samples for a shorter time span, an unavoidable consequence of frequent extinctions, were excluded from these analyses because they would be uninformative.

### SFSs in subpopulations of different ages

To test our assumption that intermediate frequency alleles are enriched after subpopulation founding, we estimated the median frequency of all minor alleles (median MAF) separately for each subpopulation sample. Then, we tested if median MAF was associated with the log_10_(age + 1) of the subpopulations using a mixed-effect model and the *lme4* v.1.1-21 (67) R package with island group of origin and subpopulation of origin (nested within island group) as random effects. For this analysis, closely neighboring islands FS and FSS, LON and LONA, and SK, SKN, SKO, and SKW were combined into island groups, as their shorelines are separated by less than 10 m (Figure 1). Statistical significance was assessed using the *lmerTest* v.3.1-3 (68) R package. Subpopulation age was calculated based on the biannual sampling data, which started in 1982 (20,22). For subpopulations that have been extant since the survey began and whose age is thus not exactly known, age was set to the duration of the survey. We tested if age groups (Young: ≤ 2 years, Intermediate: > 2 and ≤ 15, and Old: > 15; following Angst et al. (24)) differed in median MAF using Welch’s *t*-tests. For multiple samples per subpopulation, we used the average of all median MAF estimates from the given subpopulation.

### Genome-wide allele frequency changes

For consecutive samples from the same subpopulations, we tested if median MAF changed over the course of the study period using mixed-effect models with island group of origin and subpopulation of origin (nested within island group) as random effects. We also tested the same model excluding subpopulations with potential gene flow plus hybrid vigor, because under hybrid vigor, we expect a sharp temporal increase in median MAF at the time of the event rather than a linear evolution. To understand how median MAF evolves in subpopulations of different ages, we additionally tested subsets of the different age groups using the same model. One subset encompassed all the young subpopulations; another included all intermediate and old subpopulations that did not differ in median MAF in the Welch’s *t*-tests. To test if genome-wide allele frequencies change differently during the planktonic phase (spring to summer) compared to the resting phase (summer to spring), we tested if the difference in median MAF from consecutive samples of the same subpopulation differed between spring and summer samples of the same year versus between summer and the next year’s spring samples using paired Wilcoxon tests in R. In cases where several consecutive observations of a single subpopulation were available, we randomly chose one to avoid pseudo-replication.

### Temporal changes in genomic diversity and genomic differentiation

Along with temporal changes in whole-genome allele frequencies, we also investigated changes in genomic diversity and differentiation. For genomic diversity, we tested if a subpopulation’s θ was stable over time using consecutive samples from the same subpopulation. Specifically, we used linear mixed-effect models to test if θ was associated with time since the first sampling and included the subpopulation of origin nested within the island group of origin as a random effect in the model. As for our analysis of median MAF, we also tested subsets of the data with the same model. To test whether the association between θ and time depends on the degree of isolation, i.e., if a potential increase in θ over time strengthens when isolation is less, we included the interaction of time and the mean log_10_ distance to the subpopulation’s two nearest neighbors (NN2) using additional models with otherwise identical parameters. Isolation has been suggested as an important predictor of genomic diversity in this metapopulation, and NN2 a useful estimator of it (24). Regarding population differentiation, we used pairwise *F*_ST_ between all samples of the same subpopulation to test if the nonlinear increase described by *F*_ST_ = ab^1/∂t^ (32), where ∂t is the time between samples, is a better fit than a simple linear model, i.e., *F*_ST_ = a + b∂t, using AIC estimation. Furthermore, to test if there is more genomic differentiation or a greater change in θ during the planktonic phase than the resting phase, we used paired Wilcoxon tests in R as described above for median MAF. For plotting, we transformed pairwise *F*_ST_ using ln((*F*_ST_ + 0.001)/(1 + (*F*_ST_ + 0.001))).

### Empirical data of single, large, stable population

To compare subpopulations in our metapopulation with a temporal sample from a single, large, more stable population, we estimated genome-wide allele frequencies for the *D. magna* population from Oud Heverlee pond near Leuven, Belgium (50°50II22.16II N, 4°39II18.16II E). We downloaded genomic sequences of 36 individual *D. magna* from GenBank (NCBI database; Bioproject ID: PRJNA344883) that stem from sediment core samples of three times twelve genotypes about nine years apart (29) and used the above described bioinformatic pipeline to map sequencing reads and variant calling, except that we used GATK v.4.1.9.0 HaplotypeCaller (69), a more powerful approach for calling variants from single genotypes. In the resulting VCF file, we used the same parameters for filtering SNPs by quality and calculated median MAF for each of the three samples using VCFtools and R.

### Simulations

We conducted numerical simulations using the forward simulation software SLiM v.4.0.1 (70) to investigate the isolated effects of evolutionary processes on genome-wide allele frequencies. These simulations were guided by a previous simulation framework that had been calibrated with parameters that reflect the biology of the focal subpopulations (24). Specifically, we adopted the reproduction model (sexual reproduction only every eighth generation, else asexually reproducing), recombination rate (1×10^−8^), carrying capacity (300), and mutation rate (1×10^−8^) generating synonymous as well as nonsynonymous mutations (DFE for nonsynonymous mutations followed a gamma distribution with mean = −0.03 and shape = 0.2; dominance coefficient = 0.1). For each setting (= scenario), we conducted 1,000 replicate simulations of 10 Mbp sequence. A first modeled scenario, the base simulation, was subpopulation founding. For that, an average of 1.5 founders (range of founders: 1 to 7), the suggested number of colonizers in this system (23,34), was taken from a population at mutation-drift equilibrium, i.e., after a burn-in of 10,000 generations. Median MAF and θ of the newly founded subpopulation were calculated before (summer) and after (spring) sexual recombination; pairwise *F*_ST_ was calculated between all consecutive spring and summer, as well as summer and spring time points. In another scenario built on the base simulation, we included bottlenecks of size n = 10 directly after sexual recombination to examine the effects of overwintering bottlenecks. A further scenario, designed to simulate a hybrid vigor event, introduced an immigrant from an independent population at mutation-drift equilibrium into the base simulation population 10 years after the population’s founding. A final simulation focused on investigating the effects of genetic drift and newly arising mutations on the temporal dynamics of median MAF. For that, we used the parameters of the base simulation and 100 replicates for each of seven different carrying capacities between 100 and 10,000. The simulated data was analyzed in the same manner as the empirical data.

## Declarations

### Ethics approval and consent to participate

Not applicable

### Consent for publication

Not applicable

### Availability of data and materials

Analysis scripts are available at https://github.com/pascalangst/Angst_etal_2024, raw data are deposited at the NCBI SRA database (BioProject IDs PRJNA862292), and the filtered VCF file is deposited at https://doi.org/10.6084/m9.figshare.24324100.

### Competing interests

The authors declare that they have no competing interests.

## Funding

This work was supported by the Swiss National Science Foundation (SNSF) (grant numbers 310030B_166677, 310030_188887 to D.E.).

## Authors’ contributions

P.A., P.D.F., and D.E. designed the study. P.A. analyzed the data. D.E., C.R.H., and F.B.-A. collected the long-term metapopulation data. D.E. supervised the long-term metapopulation data collection and curated the data. P.D.F., D.E., and F.B.-A. collected, and P.A. and P.D.F. prepared the metapopulation samples. P.A. wrote the manuscript. All authors reviewed and approved the final manuscript.

## Acknowledgments

We thank V. Ilmari Pajunen, I. Pajunen, J. Hottinger, A. Cabalzar, M. Lehto, J. Lohr, D. Preiswerk, D. Duneau, K. I. Ebert, G. G. Ebert, A. Marcelino, Y. Haag, E. Haag, C. Liautard-Haag, C. Reisser, C. Molinier, E. Hürlimann, and C. Mills for help in the field. We thank members of the Ebert group for feedback on the study and the manuscript. Suzanne Zweizig improved the language of the text.

## Notes

### Competing Interest Statement

The authors have declared no competing interest.

