## Supplemental Material for "Genome-wide allele frequency changes reveal that dynamic metapopulations evolve differently"

^†^ Shared last author

**Figure S1: The median MAF is higher in younger subpopulations than in older subpopulations**. Estimates of samples taken from the same subpopulation are averaged and their age is log_10_(age + 1)-transformed. Colors indicate island (group*) of origin (see legend).

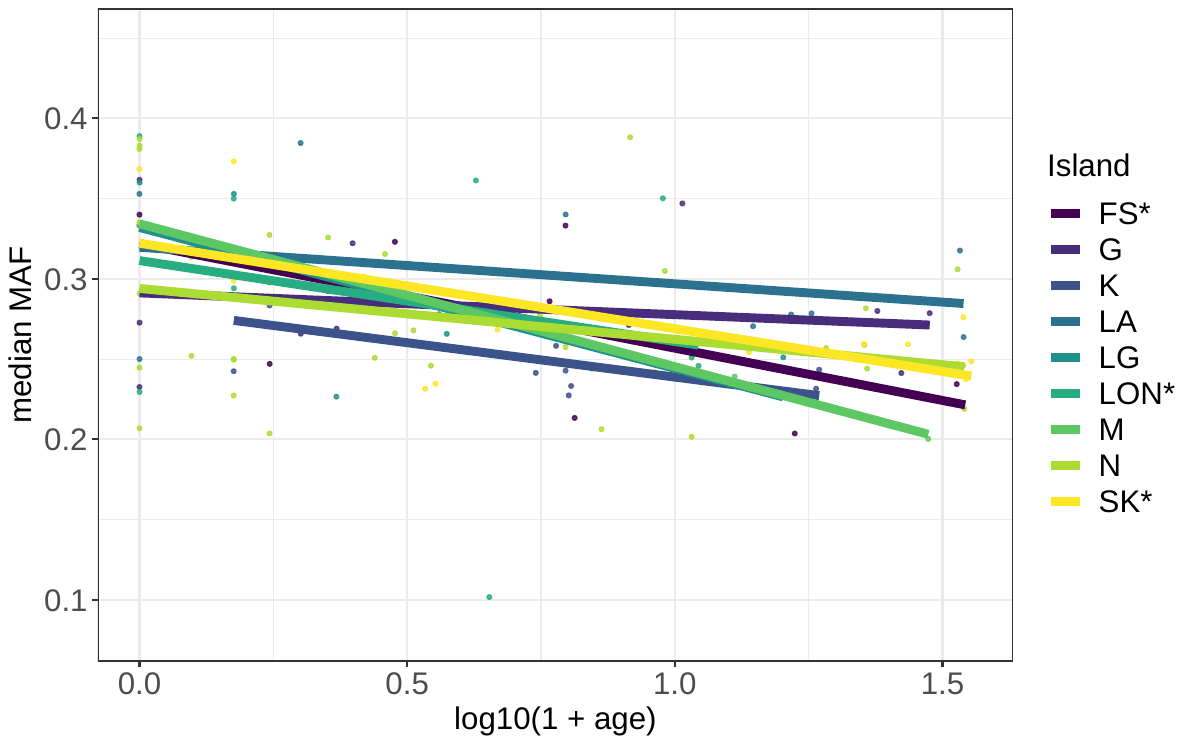

**Figure S2: Delta median MAF and Pairwise *F*_ST_** **between seasons and years per subpopulation.** (A) Median MAF is slightly increasing on average between samples of the same year, which is different from summer and consecutive spring samples, in between which median MAF decreases. (B) Pairwise *F*_ST_ between samples of consecutive years and between summer and consecutive spring samples is larger than between samples of the same year. Overall statistics derive from Kruskal-Wallis tests and individual groups were compared with paired Wilcoxon tests. The numbers below the boxes indicate (un)transformed means.

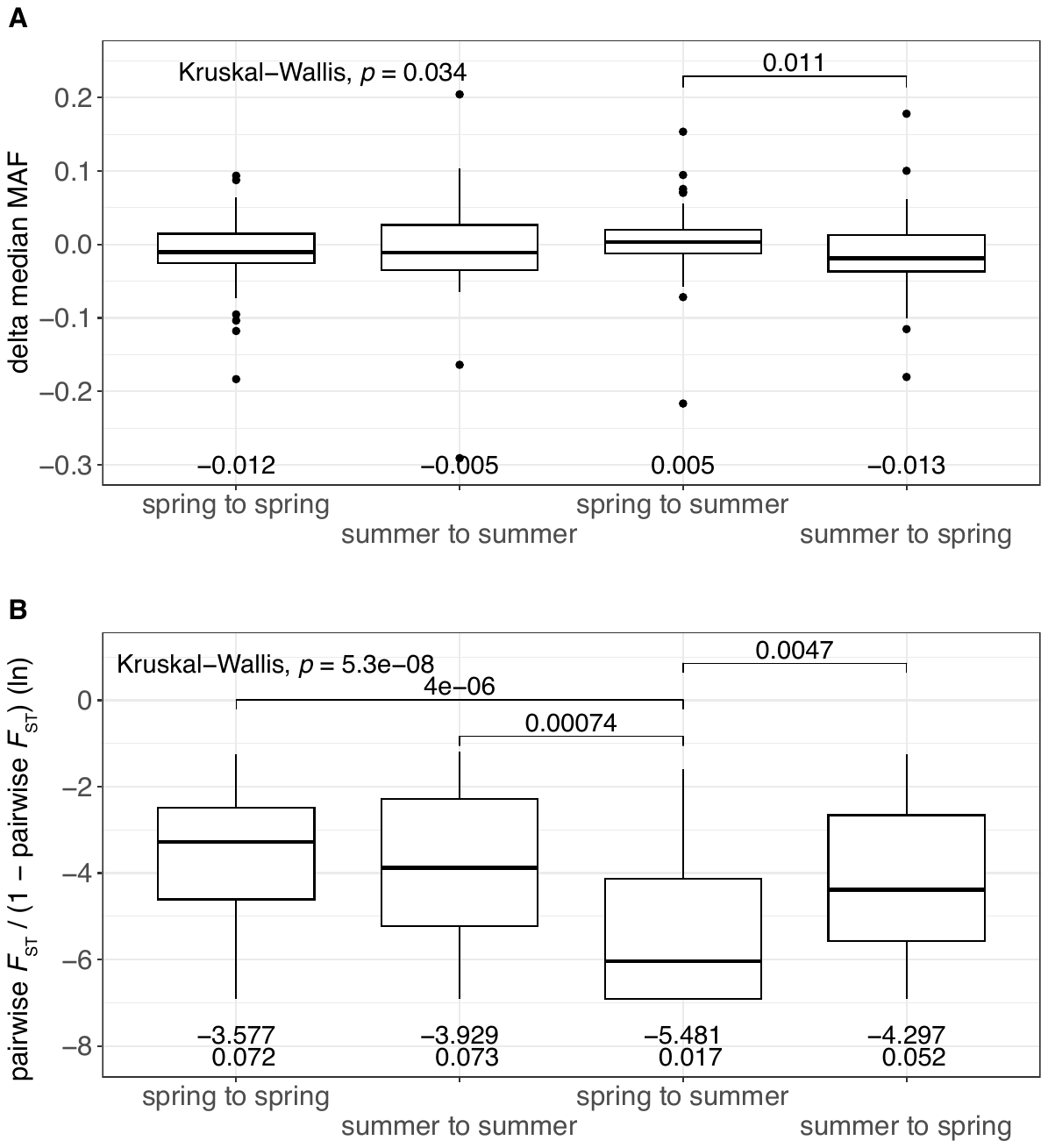

**Figure S3: Genomic diversity (θ) in the different subpopulations.** θ of individual subpopulations remains approximately stable overall during our study. Colors indicate island (group) of origin (see legend).

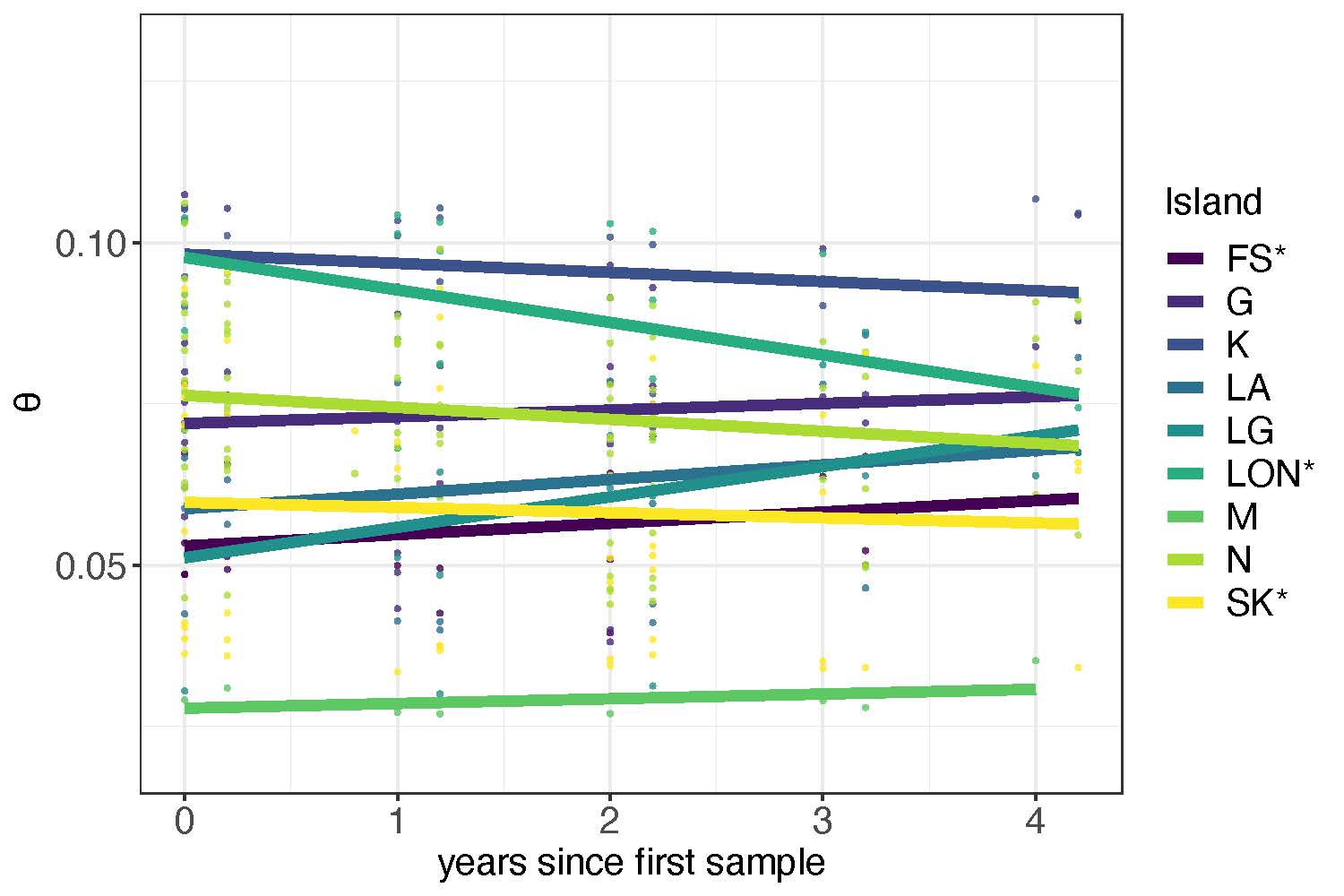

**Figure S4: Delta θ between seasons and years per subpopulation.** Changes between all possible combinations of consecutive samples do not differ. Statistics derive from a Kruskal-Wallis test.

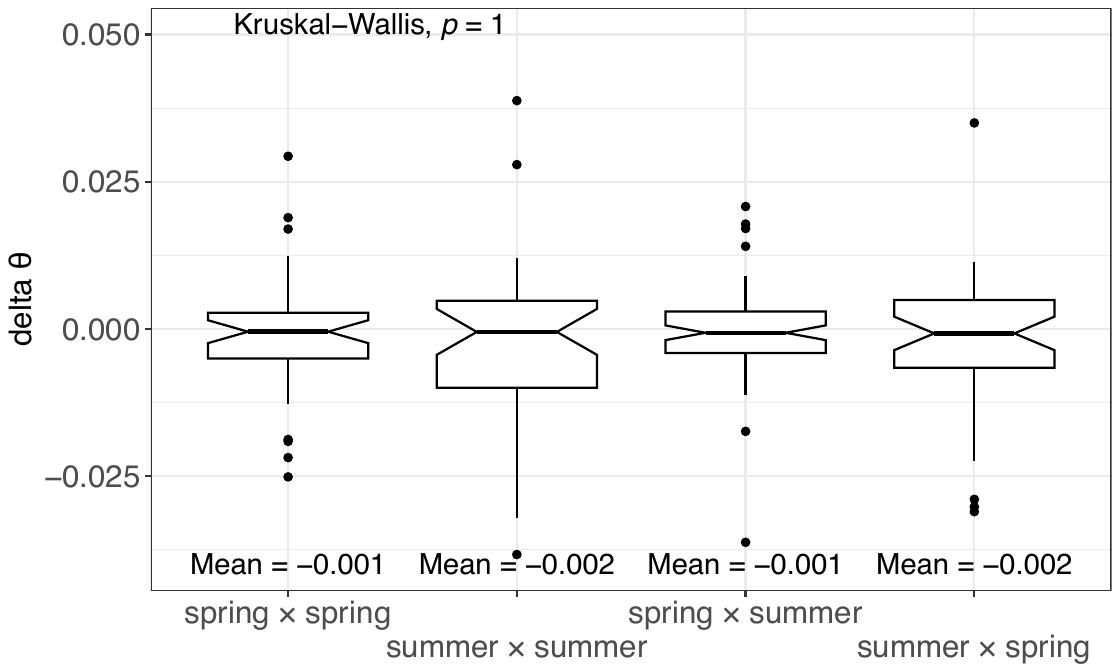

**Figure S5: Genome-wide differentiation among all samples, measured as pairwise *F*_ST_.** Pairwise *F*_ST_ heat map of *D. magna* subpopulations, darker blues indicate higher genomic differentiation and lighter blues lower differentiation. Samples are sorted alphanumerically and those from the same subpopulation and island form clusters of low differentiation.

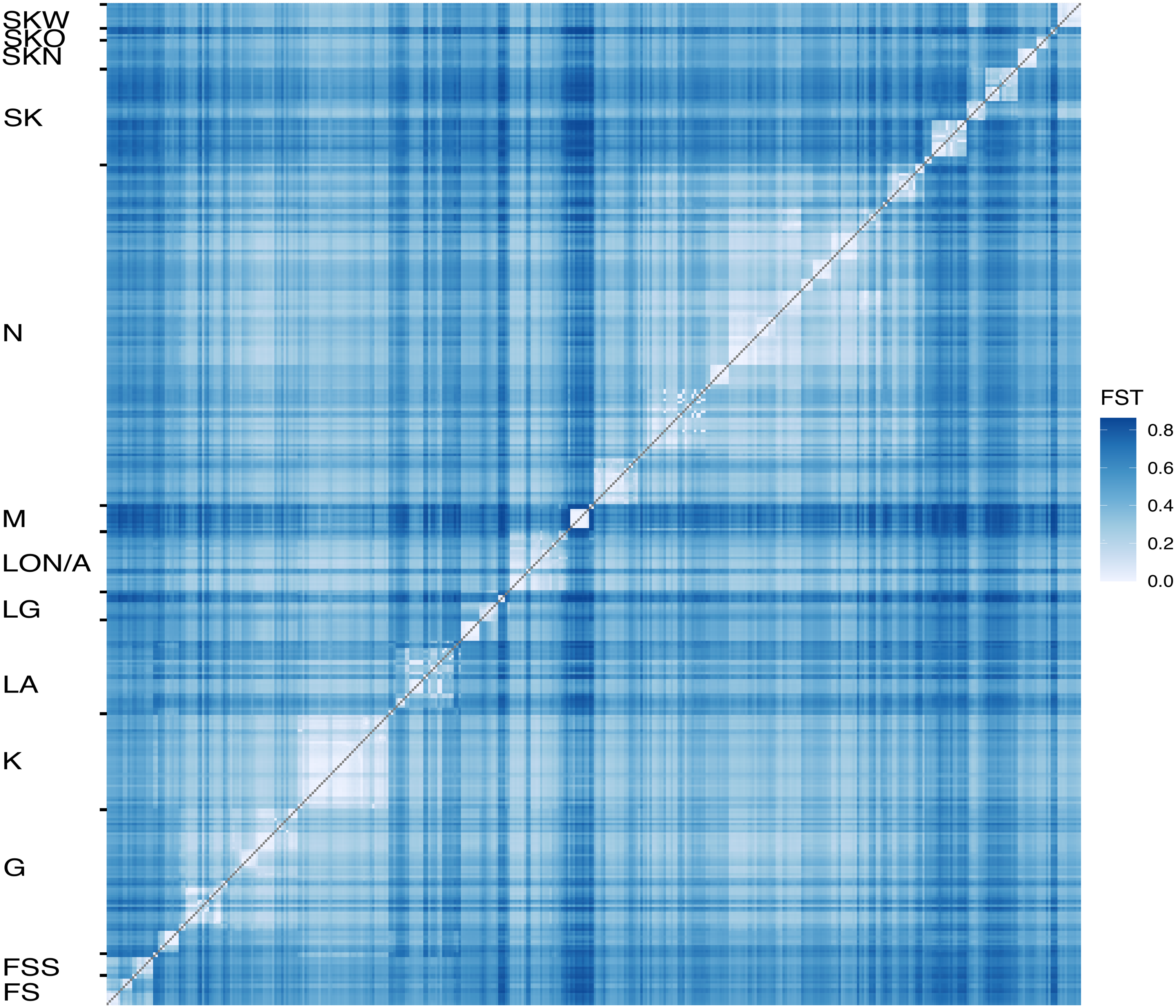

**Figure S6: Simulation to understand the effects of genetic drift and newly arising mutations on the temporal dynamics of median MAF by using different population sizes.** Using parameters of the base simulation with different carrying capacities, we observe initially more stable median MAF at intermediate frequencies in bigger populations than in smaller populations. After that, the bigger populations approach lower values over time than the smaller populations.

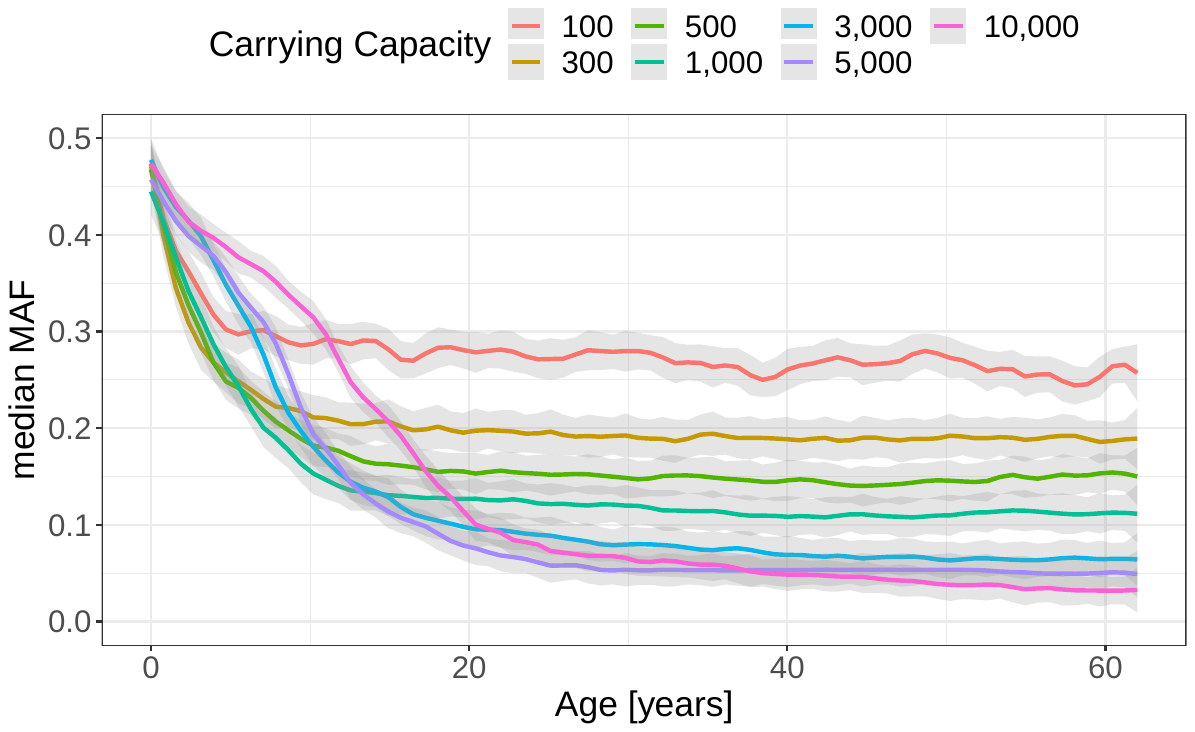

**Figure S7: Dimensionality-reduction using t-SNE.** T-SNE based on whole-genome allele frequency data reveals spatial population structure. Samples from the same island and pond form clusters in most cases. Colored symbols indicate island and pond of origin (see legend).

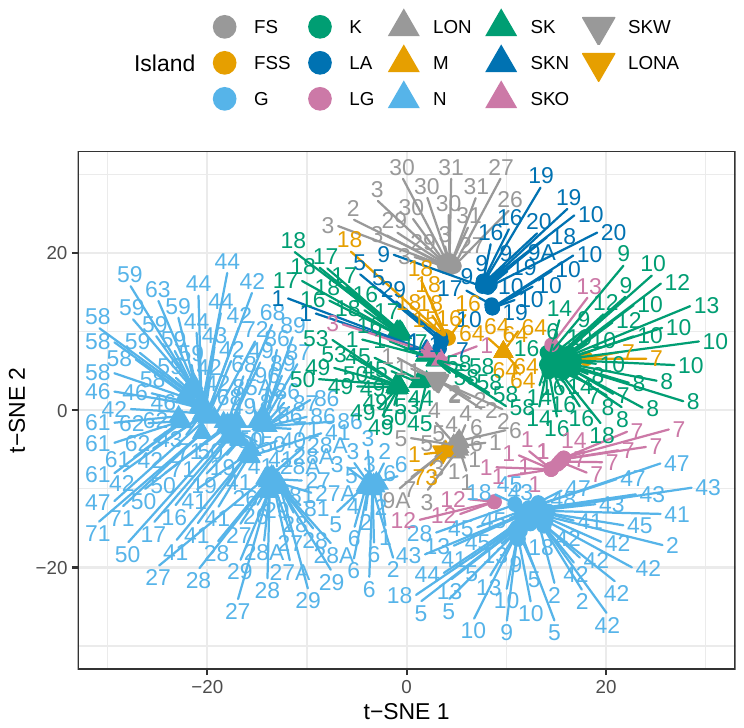

**Table S1: Detailed sample information and mapping statistics.** Each sample name (ID) consists of the pond name and the sampling time and is accompanied by the number of pooled animals prior to DNA extraction (#animals), the percentage of sequencing reads that mapped to the *D. magna* reference genome (mapping_percent), the average whole-genome coverage (average_coverage), and the infection status with *H. tvaerminnensis* (infection_status).

| ID | #animals | mapping_percent | average_coverage | infection_status |
| --- | --- | --- | --- | --- |
| FS-2_spr2014 | 9 | 0.9886 | 84.383 | 0 |
| FS-26_smr2017 | 50 | 0.9545 | 72.1964 | 0 |
| FS-27_smr2014 | 50 | 0.8854 | 21.8786 | 1 |
| FS-27_smr2018 | 50 | 0.9464 | 62.3466 | 0 |
| FS-27_spr2014 | 27 | 0.9399 | 22.6618 | 1 |
| FS-27_spr2016 | 19 | 0.9938 | 114.52 | 1 |
| FS-29_smr2014 | 50 | 0.9909 | 67.6495 | 0 |
| FS-29_spr2014 | 17 | 0.9925 | 50.4757 | 0 |
| FS-3_smr2015 | 16 | 0.9704 | 60.0981 | 1 |
| FS-3_smr2018 | 50 | 0.9904 | 49.8351 | 0 |
| FS-3_spr2014 | 11 | 0.9649 | 67.5036 | 1 |
| FS-3_spr2015 | 29 | 0.9769 | 59.31 | 1 |
| FS-3_spr2016 | 50 | 0.9797 | 75.2732 | 1 |
| FS-30_smr2015 | 44 | 0.9803 | 71.2947 | 0 |
| FS-30_smr2016 | 45 | 0.9699 | 48.9934 | 0 |
| FS-30_smr2018 | 50 | 0.9854 | 112.682 | 0 |
| FS-30_spr2015 | 40 | 0.9798 | 61.3987 | 0 |
| FS-31_smr2014 | 50 | 0.8518 | 52.3117 | 1 |
| FS-31_smr2017 | 32 | 0.5408 | 69.8623 | 1 |
| FS-31_spr2014 | 31 | 0.918 | 53.9897 | 1 |
| FSS-15_smr2015 | 41 | 0.9167 | 77.7063 | 1 |
| FSS-16_smr2016 | 50 | 0.9937 | 71.8726 | 0 |
| FSS-16_spr2016 | 50 | 0.9926 | 100.504 | 0 |
| FSS-18_smr2014 | 50 | 0.9682 | 21.7951 | 1 |
| FSS-18_smr2017 | 48 | 0.9499 | 44.725 | 1 |
| FSS-18_smr2018 | 50 | 0.9458 | 57.5265 | 1 |
| FSS-18_spr2014 | 52 | 0.9886 | 76.6795 | 1 |
| FSS-18_spr2016 | 50 | 0.928 | 80.5865 | NA |
| FSS-18_spr2017 | 50 | 0.9908 | 200.754 | NA |
| FSS-7_smr2014 | 50 | 0.8843 | 63.4516 | 1 |
| FSS-7_spr2014 | 51 | 0.8943 | 68.2905 | 1 |
| G-10_smr2016 | 48 | 0.9892 | 74.6709 | 0 |
| G-10_smr2017 | 50 | 0.7601 | 44.6196 | 0 |
| G-10_smr2018 | 50 | 0.9114 | 116.049 | 0 |
| G-13_smr2014 | 50 | 0.833 | 56.9506 | 1 |
| G-13_smr2016 | 50 | 0.8947 | 75.8125 | 1 |
| G-13_smr2017 | 50 | 0.9834 | 106.706 | 1 |
| G-13_spr2014 | 38 | 0.8331 | 19.9154 | 1 |
| G-13_spr2016 | 49 | 0.8919 | 42.7843 | 1 |
| G-18_smr2016 | 50 | 0.9776 | 83.316 | 0 |
| G-18_spr2014 | 51 | 0.9872 | 66.0887 | 0 |
| G-18_spr2016 | 49 | 0.9903 | 95.9504 | 0 |
| G-18_spr2018 | 42 | 0.9652 | 44.8112 | 0 |
| G-2_smr2015 | 50 | 0.5816 | 44.3322 | 1 |
| G-2_spr2014 | 19 | 0.9721 | 62.5149 | 0 |
| G-2_spr2015 | 23 | 0.9117 | 54.7689 | 1 |
| G-28_smr2016 | 50 | 0.9465 | 66.6268 | 1 |
| G-33_spr2014 | 39 | 0.9799 | 0.321314 | NA |
| G-41_smr2017 | 50 | 0.9905 | 90.1215 | 0 |
| G-41_smr2018 | 41 | 0.9706 | 75.6919 | 0 |
| G-41_spr2016 | 45 | 0.9882 | 66.5529 | 0 |
| G-42_smr2015 | 50 | 0.9386 | 56.8964 | 1 |
| G-42_smr2017 | 50 | 0.8642 | 185.039 | 1 |
| G-42_smr2018 | 48 | 0.4484 | 24.3485 | 1 |
| G-42_spr2014 | 51 | 0.9838 | 56.1439 | 1 |
| G-42_spr2015 | 18 | 0.9924 | 62.3006 | 1 |
| G-42_spr2016 | 50 | 0.8958 | 45.9702 | 1 |
| G-42_spr2017 | 50 | 0.9731 | 49.5918 | 1 |
| G-42_spr2018 | 50 | 0.5695 | 36.2327 | 1 |
| G-43_smr2014 | 50 | 0.845 | 20.3192 | 1 |
| G-43_smr2015 | 50 | 0.8471 | 70.311 | 1 |
| G-43_smr2016 | 47 | 0.7951 | 62.1451 | 1 |
| G-43_smr2018 | 46 | 0.908 | 145.354 | 1 |
| G-43_spr2014 | 50 | 0.9507 | 23.5622 | 1 |
| G-43_spr2016 | 50 | 0.8453 | 53.1574 | 1 |
| G-43_spr2017 | 50 | 0.8218 | 40.1188 | 1 |
| G-44_spr2017 | 50 | 0.9905 | 94.6468 | 0 |
| G-45_smr2014 | 50 | 0.9819 | 53.6246 | 0 |
| G-45_smr2017 | 40 | 0.9875 | 162.166 | 0 |
| G-45_spr2014 | 44 | 0.9876 | 55.301 | 0 |
| G-45_spr2015 | 25 | 0.9893 | 56.5611 | 0 |
| G-45_spr2016 | 50 | 0.9902 | 108.814 | 0 |
| G-47_smr2017 | 50 | 0.9749 | 94.6466 | 0 |
| G-47_smr2018 | 49 | 0.9547 | 56.7243 | 1 |
| G-47_spr2018 | 50 | 0.8148 | 52.2292 | 1 |
| G-48_smr2017 | 50 | 0.9892 | 48.134 | 0 |
| G-5_smr2016 | 50 | 0.9879 | 103.563 | 1 |
| G-5_smr2017 | 50 | 0.9754 | 119.986 | 1 |
| G-5_smr2018 | 49 | 0.9001 | 54.6514 | 1 |
| G-5_spr2016 | 45 | 0.9266 | 72.424 | 1 |
| G-5_spr2017 | 39 | 0.9922 | 79.3327 | 1 |
| G-9_smr2017 | 32 | 0.991 | 48.8768 | 0 |
| G-9_smr2018 | 46 | 0.9865 | 56.0549 | 0 |
| K-10_smr2014 | 50 | 0.8134 | 50.4775 | 1 |
| K-10_smr2015 | 50 | 0.7337 | 48.8318 | 1 |
| K-10_smr2016 | 45 | 0.8627 | 80.5784 | 1 |
| K-10_smr2018 | 50 | 0.9536 | 60.039 | 1 |
| K-10_spr2014 | 28 | 0.8706 | 45.5364 | 1 |
| K-10_spr2015 | 50 | 0.9501 | 63.7988 | 1 |
| K-10_spr2016 | 45 | 0.8418 | 53.1716 | 1 |
| K-10_spr2017 | 30 | 0.9628 | 83.323 | 1 |
| K-10_spr2018 | 25 | 0.9065 | 47.4856 | 1 |
| K-12_smr2015 | 25 | 0.771 | 69.8134 | 1 |
| K-12_smr2016 | 45 | 0.8503 | 85.4343 | 1 |
| K-12_spr2015 | 50 | 0.9202 | 64.1778 | 1 |
| K-13_spr2016 | 29 | 0.9227 | 58.3655 | 1 |
| K-14_smr2015 | 50 | 0.9215 | 84.0098 | 1 |
| K-14_smr2016 | 50 | 0.9822 | 98.0483 | 1 |
| K-14_spr2014 | 66 | 0.9432 | 48.357 | 1 |
| K-14_spr2015 | 10 | 0.9077 | 53.6066 | 1 |
| K-15_spr2014 | 67 | 0.8737 | 30.4633 | 1 |
| K-16_smr2015 | 50 | 0.9237 | 62.5458 | 1 |
| K-16_smr2016 | 50 | 0.9913 | 110.594 | 1 |
| K-16_spr2014 | 54 | 0.8821 | 59.2523 | 1 |
| K-16_spr2015 | 41 | 0.9402 | 58.362 | 1 |
| K-16_spr2016 | 19 | 0.9643 | 66.8055 | 1 |
| K-18_spr2014 | 61 | 0.8831 | 64.912 | 1 |
| K-6_smr2014 | 50 | 0.9254 | 51.3405 | 1 |
| K-6_spr2014 | 53 | 0.9642 | 21.4087 | 1 |
| K-7_smr2014 | 50 | 0.9655 | 19.6351 | 1 |
| K-7_spr2014 | 71 | 0.9766 | 104.367 | 1 |
| K-7_spr2015 | 50 | 0.9915 | 65.8317 | 1 |
| K-8_smr2015 | 50 | 0.6121 | 49.2575 | 1 |
| K-8_smr2016 | 35 | 0.7786 | 59.6002 | 1 |
| K-8_smr2018 | 50 | 0.8206 | 51.2137 | 1 |
| K-8_spr2014 | 53 | 0.8848 | 62.1974 | 1 |
| K-8_spr2015 | 50 | 0.8534 | 53.026 | 1 |
| K-8_spr2017 | 50 | 0.9247 | 159.947 | 1 |
| K-8_spr2018 | 46 | 0.8335 | 42.9603 | 1 |
| K-9_smr2015 | 50 | 0.9877 | 174.801 | 1 |
| K-9_spr2014 | 58 | 0.9571 | 50.9035 | 1 |
| K-9_spr2015 | 49 | 0.9583 | 373.621 | 1 |
| LA-10_smr2014 | 50 | 0.9755 | 48.9168 | 1 |
| LA-10_smr2016 | 50 | 0.9519 | 106.391 | 1 |
| LA-10_smr2017 | 50 | 0.9695 | 46.87 | 1 |
| LA-10_smr2018 | 29 | 0.9816 | 55.7225 | 1 |
| LA-10_spr2014 | 28 | 0.9823 | 21.1503 | 1 |
| LA-10_spr2015 | 50 | 0.9808 | 50.5292 | 1 |
| LA-10_spr2016 | 60 | 0.9543 | 67.3733 | 1 |
| LA-10_spr2017 | 50 | 0.9801 | 90.7836 | 1 |
| LA-16_smr2015 | 50 | 0.6594 | 47.1419 | 1 |
| LA-16_spr2015 | 24 | 0.9608 | 58.5965 | 1 |
| LA-17_smr2018 | 50 | 0.9448 | 63.4971 | 0 |
| LA-18_smr2014 | 50 | 0.9856 | 22.5557 | 1 |
| LA-18_smr2016 | 50 | 0.9919 | 66.7712 | 1 |
| LA-18_spr2014 | 30 | 0.9877 | 20.8245 | 1 |
| LA-19_smr2014 | 50 | 0.9796 | 109.855 | 1 |
| LA-19_smr2015 | 50 | 0.951 | 55.9323 | 1 |
| LA-19_smr2016 | 50 | 0.9917 | 87.1379 | 1 |
| LA-19_smr2017 | 50 | 0.973 | 54.8301 | 1 |
| LA-19_spr2014 | 71 | 0.9845 | 21.7759 | 1 |
| LA-20_smr2015 | 50 | 0.8591 | 73.961 | 1 |
| LA-20_smr2016 | 49 | 0.99 | 77.2345 | 1 |
| LA-29_smr2018 | 50 | 0.9895 | 79.3881 | 0 |
| LA-29_spr2014 | 59 | 0.9804 | 24.0561 | 0 |
| LA-5_smr2015 | 50 | 0.9113 | 64.3915 | 0 |
| LA-5_spr2015 | 50 | 0.9871 | 60.1637 | 0 |
| LA-7_smr2017 | 50 | 0.9134 | 42.9593 | 0 |
| LA-9_smr2017 | 50 | 0.9605 | 79.0853 | 0 |
| LA-9_smr2018 | 48 | 0.9718 | 46.1944 | 0 |
| LA-9_spr2016 | 60 | 0.9347 | 47.5133 | 0 |
| LA-9_spr2017 | 50 | 0.9917 | 73.7815 | 0 |
| LA-9A_smr2018 | 29 | 0.9906 | 60.7582 | 0 |
| LG-1_smr2015 | 50 | 0.987 | 53.5693 | 1 |
| LG-1_smr2017 | 50 | 0.9731 | 67.5503 | 1 |
| LG-1_smr2018 | 50 | 0.9674 | 52.4829 | 1 |
| LG-1_spr2014 | 40 | 0.9873 | 64.8407 | 1 |
| LG-1_spr2015 | 20 | 0.9916 | 57.7477 | 1 |
| LG-1_spr2016 | 60 | 0.9575 | 81.5514 | 1 |
| LG-1_spr2017 | 50 | 0.9738 | 51.2383 | 1 |
| LG-1_spr2018 | 50 | 0.9482 | 53.794 | 1 |
| LG-12_smr2017 | 50 | 0.9298 | 43.0491 | 0 |
| LG-12_smr2018 | 46 | 0.985 | 72.6489 | 0 |
| LG-12_spr2016 | 50 | 0.9724 | 76.7427 | 0 |
| LG-13_spr2015 | 22 | 0.9861 | 54.5154 | 0 |
| LG-14_smr2018 | 50 | 0.9773 | 48.2999 | 0 |
| LG-7_smr2015 | 50 | 0.9926 | 102.572 | 0 |
| LG-7_smr2016 | 50 | 0.9867 | 100.666 | 0 |
| LG-7_smr2017 | 29 | 0.6989 | 34.5469 | 0 |
| LG-7_smr2018 | 50 | 0.926 | 65.0203 | 0 |
| LG-7_spr2014 | 49 | 0.9889 | 43.4276 | 0 |
| LG-7_spr2015 | 24 | 0.99 | 50.4875 | 0 |
| LG-7_spr2016 | 50 | 0.9912 | 93.7667 | 0 |
| LG-7_spr2017 | 38 | 0.9871 | 47.3222 | 0 |
| LON-1_smr2014 | 50 | 0.6515 | 13.7111 | 1 |
| LON-1_smr2015 | 50 | 0.6328 | 38.6659 | 1 |
| LON-1_smr2016 | 50 | 0.4055 | 30.9359 | 1 |
| LON-1_smr2018 | 50 | 0.9577 | 57.2 | 1 |
| LON-1_spr2014 | 58 | 0.8055 | 37.0342 | 1 |
| LON-1_spr2015 | 50 | 0.7053 | 44.493 | 1 |
| LON-1_spr2016 | 60 | 0.7627 | 50.5315 | 1 |
| LON-1_spr2017 | 50 | 0.8373 | 99.5763 | 1 |
| LON-3_smr2014 | 50 | 0.8814 | 64.5399 | 1 |
| LON-3_spr2014 | 63 | 0.8924 | 62.8305 | 1 |
| LON-4_smr2015 | 50 | 0.5307 | 43.4491 | 1 |
| LON-4_smr2016 | 50 | 0.899 | 64.7932 | 1 |
| LON-4_spr2014 | 55 | 0.7394 | 31.6327 | 1 |
| LON-4_spr2015 | 50 | 0.8167 | 45.7243 | 1 |
| LON-5_smr2015 | 50 | 0.78 | 66.9466 | 1 |
| LON-5_smr2016 | 50 | 0.9495 | 73.2112 | 1 |
| LON-5_spr2014 | 53 | 0.9572 | 48.7995 | 1 |
| LON-5_spr2015 | 50 | 0.956 | 62.4419 | 1 |
| LON-6_smr2016 | 50 | 0.9458 | 92.3743 | 1 |
| LON-6_smr2018 | 50 | 0.8453 | 79.2169 | 1 |
| LON-6_spr2014 | 51 | 0.9094 | 20.9864 | 1 |
| LON-6_spr2016 | 45 | 0.7895 | 58.9687 | 1 |
| LON-7_smr2018 | 50 | 0.9422 | 101.797 | 1 |
| LON-9A_smr2018 | 50 | 0.8964 | 52.7198 | 0 |
| LON-9A_spr2014 | 14 | 0.955 | 25.4082 | 1 |
| LONA-1_smr2014 | 50 | 0.9667 | 92.5223 | 1 |
| M-62_spr2014 | 50 | 0.948 | 24.7763 | 1 |
| M-64_smr2014 | 50 | 0.9325 | 20.0168 | 0 |
| M-64_smr2015 | 50 | 0.9206 | 59.199 | 0 |
| M-64_smr2017 | 47 | 0.9872 | 48.0499 | 0 |
| M-64_spr2014 | 42 | 0.9871 | 47.523 | 0 |
| M-64_spr2015 | 23 | 0.9924 | 84.6306 | 0 |
| M-64_spr2016 | 46 | 0.9536 | 53.8583 | 0 |
| M-64_spr2017 | 50 | 0.9915 | 123.482 | 0 |
| M-64_spr2018 | 48 | 0.9838 | 50.1236 | 0 |
| M-73_smr2014 | 50 | 0.9641 | 79.7032 | 1 |
| M-73_spr2014 | 20 | 0.9679 | 110.979 | 1 |
| N-1_spr2014 | 11 | 0.9729 | 46.767 | 1 |
| N-16_smr2016 | 50 | 0.9911 | 90.6979 | 0 |
| N-17_smr2016 | 50 | 0.9821 | 54.5953 | 0 |
| N-19_smr2018 | 43 | 0.9868 | 57.7156 | 1 |
| N-19_spr2018 | 25 | 0.8629 | 80.9195 | 1 |
| N-2_smr2015 | 50 | 0.8002 | 42.7695 | 1 |
| N-2_smr2016 | 37 | 0.9514 | 46.7071 | 1 |
| N-2_spr2015 | 51 | 0.9549 | 55.1849 | 1 |
| N-2_spr2016 | 60 | 0.9175 | 51.2819 | 1 |
| N-26_smr2018 | 50 | 0.9672 | 60.6573 | 1 |
| N-26_spr2018 | 46 | 0.9434 | 56.6389 | 1 |
| N-27_smr2014 | 50 | 0.9319 | 43.4323 | 1 |
| N-27_smr2015 | 50 | 0.6773 | 43.9352 | 1 |
| N-27_smr2018 | 26 | 0.8173 | 63.0738 | 1 |
| N-27_spr2014 | 49 | 0.9625 | 20.2174 | 1 |
| N-27_spr2015 | 50 | 0.9677 | 59.5057 | 1 |
| N-27_spr2016 | 50 | 0.8238 | 51.4182 | 1 |
| N-27A_smr2014 | 50 | 0.6979 | 39.5103 | 1 |
| N-27A_spr2014 | 51 | 0.834 | 19.1585 | 1 |
| N-27A_spr2015 | 50 | 0.9517 | 125.623 | 1 |
| N-28_smr2014 | 50 | 0.8883 | 56.7589 | 1 |
| N-28_smr2016 | 50 | 0.9821 | 126.296 | 1 |
| N-28_smr2018 | 50 | 0.787 | 37.3607 | 1 |
| N-28_spr2014 | 49 | 0.9033 | 59.5271 | 1 |
| N-28_spr2016 | 31 | 0.9039 | 54.9076 | 1 |
| N-28A_smr2014 | 50 | 0.713 | 38.0498 | 1 |
| N-28A_smr2015 | 44 | 0.8391 | 76.7967 | 1 |
| N-28A_smr2016 | 50 | 0.9921 | 90.5412 | 1 |
| N-28A_smr2018 | 50 | 0.803 | 45.0244 | 1 |
| N-28A_spr2015 | 43 | 0.9427 | 67.6593 | 1 |
| N-29_smr2016 | 50 | 0.9915 | 60.0775 | 1 |
| N-29_smr2018 | 50 | 0.9437 | 63.5656 | 1 |
| N-29_spr2016 | 12 | 0.9935 | 54.8293 | 1 |
| N-29_spr2018 | 41 | 0.9733 | 46.0413 | 1 |
| N-3_smr2014 | 50 | 0.8895 | 35.5685 | 1 |
| N-3_smr2015 | 50 | 0.8038 | 60.2704 | 1 |
| N-3_smr2016 | 50 | 0.8674 | 94.9034 | 1 |
| N-3_spr2014 | 56 | 0.931 | 18.6522 | 1 |
| N-3_spr2015 | 48 | 0.8984 | 68.1039 | 1 |
| N-4_smr2016 | 50 | 0.9223 | 63.6505 | 1 |
| N-40_smr2014 | 50 | 0.7839 | 36.424 | 1 |
| N-40_spr2014 | 46 | 0.8756 | 18.6395 | 1 |
| N-41_smr2014 | 50 | 0.9906 | 51.865 | 1 |
| N-41_smr2015 | 50 | 0.8991 | 101.942 | 1 |
| N-41_smr2016 | 50 | 0.787 | 62.6386 | 1 |
| N-41_smr2017 | 50 | 0.5324 | 75.5459 | 1 |
| N-41_spr2014 | 49 | 0.9925 | 61.2961 | 0 |
| N-41_spr2015 | 50 | 0.9886 | 51.468 | 1 |
| N-41_spr2016 | 50 | 0.7721 | 42.2087 | 1 |
| N-41_spr2017 | 50 | 0.7946 | 83.851 | 1 |
| N-42_smr2014 | 50 | 0.9269 | 37.1563 | 1 |
| N-42_smr2015 | 50 | 0.9483 | 75.6429 | 1 |
| N-42_smr2016 | 50 | 0.9689 | 60.2848 | 1 |
| N-42_smr2017 | 50 | 0.9462 | 97.8003 | 1 |
| N-42_spr2014 | 48 | 0.95 | 108.054 | 1 |
| N-42_spr2015 | 50 | 0.9827 | 51.9164 | 1 |
| N-42_spr2016 | 50 | 0.9921 | 89.0543 | 1 |
| N-42_spr2017 | 33 | 0.9914 | 44.6437 | 1 |
| N-43_smr2014 | 50 | 0.8337 | 27.7869 | 1 |
| N-43_smr2016 | 50 | 0.9758 | 86.8556 | 1 |
| N-43_spr2014 | 53 | 0.9034 | 16.1057 | 1 |
| N-43_spr2016 | 29 | 0.9198 | 81.7096 | 1 |
| N-44_smr2014 | 50 | 0.9114 | 38.7979 | 1 |
| N-44_smr2015 | 50 | 0.6471 | 48.9093 | 1 |
| N-44_smr2016 | 50 | 0.83 | 92.3578 | 1 |
| N-44_spr2014 | 49 | 0.938 | 23.3142 | 1 |
| N-44_spr2015 | 50 | 0.8516 | 57.396 | 1 |
| N-46_smr2017 | 50 | 0.738 | 126.874 | 1 |
| N-46_smr2018 | 46 | 0.8026 | 44.1801 | 1 |
| N-46_spr2018 | 50 | 0.6353 | 68.2285 | 1 |
| N-47_spr2016 | 52 | 0.9671 | 63.1052 | 1 |
| N-49_smr2017 | 50 | 0.9869 | 110.236 | 1 |
| N-49_spr2016 | 54 | 0.9261 | 58.3954 | 1 |
| N-5_smr2016 | 50 | 0.816 | 73.3776 | 1 |
| N-5_spr2016 | 56 | 0.8313 | 58.3995 | 1 |
| N-50_smr2014 | 50 | 0.9026 | 41.8904 | 1 |
| N-50_smr2016 | 50 | 0.8792 | 81.7859 | 1 |
| N-50_smr2017 | 50 | 0.9825 | 115.957 | 1 |
| N-50_smr2018 | 42 | 0.5465 | 48.1678 | 1 |
| N-50_spr2014 | 54 | 0.9432 | 21.2262 | 1 |
| N-50_spr2015 | 50 | 0.948 | 87.6539 | 1 |
| N-50_spr2016 | 65 | 0.8289 | 47.2191 | 1 |
| N-50_spr2018 | 41 | 0.746 | 44.4829 | 1 |
| N-58_smr2015 | 50 | 0.8149 | 71.5684 | 1 |
| N-58_smr2016 | 50 | 0.9073 | 66.3786 | 1 |
| N-58_smr2018 | 45 | 0.8649 | 57.6587 | 1 |
| N-58_spr2015 | 42 | 0.9779 | 50.7495 | 1 |
| N-58_spr2016 | 80 | 0.8066 | 58.5243 | 1 |
| N-59_smr2014 | 50 | 0.7664 | 52.7019 | 1 |
| N-59_smr2015 | 50 | 0.5343 | 47.7083 | 1 |
| N-59_smr2016 | 50 | 0.7453 | 61.8021 | 1 |
| N-59_smr2018 | 50 | 0.6877 | 57.6962 | 1 |
| N-59_spr2014 | 36 | 0.8319 | 54.2554 | 1 |
| N-59_spr2015 | 50 | 0.8498 | 73.6658 | 1 |
| N-59_spr2016 | 50 | 0.633 | 41.7165 | 1 |
| N-59_spr2017 | 50 | 0.9383 | 49.6017 | 1 |
| N-6_smr2014 | 50 | 0.9302 | 59.7525 | 1 |
| N-6_smr2016 | 50 | 0.9779 | 73.0588 | 1 |
| N-6_smr2017 | 20 | 0.8915 | 85.721 | 1 |
| N-6_spr2014 | 53 | 0.9571 | 36.5148 | 1 |
| N-6_spr2016 | 51 | 0.9917 | 77.7559 | 1 |
| N-6_spr2018 | 50 | 0.9905 | 96.7448 | 1 |
| N-61_smr2014 | 50 | 0.5164 | 12.7383 | 1 |
| N-61_smr2015 | 20 | 0.4914 | 29.1687 | 1 |
| N-61_smr2016 | 27 | 0.8417 | 69.1091 | 1 |
| N-61_smr2017 | 20 | 0.7219 | 34.1326 | 1 |
| N-61_smr2018 | 50 | 0.8754 | 78.3465 | 1 |
| N-61_spr2014 | 37 | 0.7418 | 17.5543 | 1 |
| N-61_spr2018 | 48 | 0.9178 | 135.054 | 1 |
| N-62_smr2014 | 50 | 0.7986 | 30.5407 | 1 |
| N-62_smr2018 | 50 | 0.8309 | 56.1288 | 1 |
| N-62_spr2014 | 57 | 0.9247 | 36.0808 | 1 |
| N-62_spr2015 | 30 | 0.7865 | 44.1071 | 1 |
| N-62_spr2016 | 16 | 0.9601 | 108.522 | 1 |
| N-63_smr2016 | 50 | 0.9865 | 54.9043 | 0 |
| N-68_smr2016 | 25 | 0.4092 | 23.0449 | 1 |
| N-68_smr2018 | 50 | 0.8639 | 47.7465 | 1 |
| N-68_spr2016 | 60 | 0.788 | 73.5058 | 1 |
| N-68_spr2018 | 38 | 0.6431 | 104.738 | 1 |
| N-71_smr2014 | 50 | 0.6002 | 31.7977 | 1 |
| N-71_smr2015 | 50 | 0.49 | 36.7324 | 1 |
| N-71_smr2018 | 50 | 0.885 | 71.2863 | 1 |
| N-71_spr2014 | 54 | 0.7249 | 15.3076 | 1 |
| N-71_spr2015 | 50 | 0.6724 | 1.21757 | 1 |
| N-71_spr2016 | 70 | 0.5671 | 55.2987 | 1 |
| N-71_spr2018 | 50 | 0.8428 | 101.502 | 1 |
| N-72_smr2017 | 50 | 0.9825 | 167.43 | 0 |
| N-81_smr2016 | 50 | 0.5568 | 35.4669 | 1 |
| N-81_spr2016 | 70 | 0.8087 | 56.135 | 1 |
| N-85A_smr2014 | 50 | 0.8274 | 35.2537 | 1 |
| N-85A_spr2014 | 54 | 0.8932 | 18.5148 | 1 |
| N-86_smr2014 | 50 | 0.9377 | 52.0479 | 1 |
| N-86_smr2016 | 50 | 0.8388 | 61.934 | 1 |
| N-86_smr2018 | 42 | 0.5646 | 37.9813 | 1 |
| N-86_spr2014 | 49 | 0.979 | 51.3107 | 1 |
| N-86_spr2015 | 19 | 0.9528 | 109.752 | 1 |
| N-86_spr2016 | 50 | 0.9419 | 82.9051 | 1 |
| N-86_spr2018 | 50 | 0.9636 | 45.6415 | 1 |
| N-87_smr2015 | 50 | 0.6696 | 53.2414 | 1 |
| N-87_smr2016 | 50 | 0.8889 | 43.3123 | 1 |
| N-87_spr2015 | 50 | 0.9166 | 65.745 | 1 |
| N-89_smr2014 | 50 | 0.5616 | 9.51041 | 1 |
| N-89_smr2015 | 50 | 0.5483 | 52.5007 | 1 |
| N-89_spr2014 | 50 | 0.8405 | 30.9543 | 1 |
| N-89_spr2015 | 50 | 0.9226 | 65.1159 | 1 |
| N-96_spr2016 | 75 | 0.77 | 52.537 | 1 |
| SK-1_smr2014 | 50 | 0.9318 | 90.1333 | 1 |
| SK-1_smr2016 | 50 | 0.9822 | 60.4223 | 1 |
| SK-1_spr2014 | 61 | 0.9458 | 0.230235 | NA |
| SK-1_spr2015 | 50 | 0.8924 | 44.5172 | 1 |
| SK-16_smr2014 | 50 | 0.8686 | 26.7496 | 1 |
| SK-16_smr2016 | 50 | 0.9775 | 383.063 | 1 |
| SK-16_smr2017 | 50 | 0.9437 | 63.1383 | 1 |
| SK-16_spr2014 | 60 | 0.9866 | 20.1261 | 0 |
| SK-16_spr2016 | 52 | 0.9699 | 62.0016 | 1 |
| SK-16_spr2017 | 50 | 0.9767 | 99.2169 | 1 |
| SK-17_smr2016 | 50 | 0.8215 | 49.7722 | 1 |
| SK-17_smr2017 | 50 | 0.9554 | 185.786 | 1 |
| SK-17_smr2018 | 50 | 0.5907 | 36.1063 | 1 |
| SK-17_spr2016 | 60 | 0.8014 | 62.3768 | 1 |
| SK-17_spr2018 | 45 | 0.964 | 49.0833 | 1 |
| SK-18_smr2015 | 50 | 0.9862 | 54.6725 | 0 |
| SK-18_smr2016 | 50 | 0.6617 | 83.9426 | 1 |
| SK-18_spr2015 | 31 | 0.9772 | 53.5491 | 0 |
| SK-18_spr2016 | 36 | 0.7129 | 40.2385 | 1 |
| SK-44_smr2017 | 44 | 0.9925 | 132.07 | 0 |
| SK-45_smr2014 | 50 | 0.7815 | 27.5791 | 1 |
| SK-45_smr2015 | 50 | 0.7721 | 50.1728 | 1 |
| SK-45_smr2016 | 50 | 0.8898 | 62.46 | 1 |
| SK-45_smr2017 | 22 | 0.931 | 83.6568 | 1 |
| SK-45_smr2018 | 50 | 0.7617 | 46.6474 | 1 |
| SK-45_spr2014 | 37 | 0.8716 | 43.4562 | 1 |
| SK-45_spr2016 | 45 | 0.9506 | 96.827 | 1 |
| SK-49_smr2015 | 50 | 0.9858 | 82.0767 | 0 |
| SK-49_smr2016 | 50 | 0.9766 | 51.7639 | 0 |
| SK-49_smr2018 | 39 | 0.9751 | 55.847 | 0 |
| SK-49_spr2014 | 50 | 0.9935 | 27.6059 | 0 |
| SK-49_spr2015 | 50 | 0.9898 | 68.8115 | 0 |
| SK-49_spr2016 | 50 | 0.9925 | 60.5854 | 0 |
| SK-49_spr2017 | 50 | 0.9761 | 89.8514 | 0 |
| SK-50_smr2014 | 50 | 0.9915 | 56.5774 | 0 |
| SK-50_smr2015 | 50 | 0.9899 | 63.1238 | 0 |
| SK-50_smr2016 | 50 | 0.9894 | 73.6538 | 0 |
| SK-50_spr2014 | 48 | 0.9888 | 47.9354 | 0 |
| SK-53_smr2015 | 50 | 0.9916 | 100.368 | 1 |
| SK-53_smr2016 | 50 | 0.9372 | 53.4103 | 1 |
| SK-53_spr2015 | 16 | 0.989 | 53.7698 | 1 |
| SK-58_smr2014 | 50 | 0.9289 | 44.2539 | 1 |
| SK-58_smr2016 | 50 | 0.5724 | 61.8007 | 1 |
| SK-58_smr2017 | 50 | 0.411 | 19.8792 | 1 |
| SK-58_smr2018 | 50 | 0.7355 | 68.4431 | 1 |
| SK-58_spr2014 | 48 | 0.9683 | 19.4414 | 1 |
| SK-58_spr2015 | 50 | 0.9437 | 136.891 | 1 |
| SK-58_spr2016 | 38 | 0.8705 | 60.2672 | 1 |
| SK-58_spr2017 | 50 | 0.777 | 36.6504 | 1 |
| SKN-1_smr2015 | 50 | 0.7585 | 46.7005 | 1 |
| SKN-1_smr2016 | 50 | 0.9142 | 51.7608 | 1 |
| SKN-1_spr2014 | 49 | 0.9006 | 52.9265 | 1 |
| SKN-1_spr2015 | 46 | 0.9498 | 77.9327 | 1 |
| SKN-2_spr2014 | 61 | 0.9744 | 21.7282 | 1 |
| SKO-1_spr2014 | 45 | 0.9706 | 57.1013 | 1 |
| SKO-3_smr2014 | 50 | 0.9883 | 37.4972 | 0 |
| SKO-3_spr2014 | 51 | 0.9883 | 37.4972 | 0 |
| SKO-3_spr2015 | 50 | 0.9871 | 63.7467 | 0 |
| SKW-1_smr2014 | 50 | 0.9867 | 61.2855 | 1 |
| SKW-1_smr2015 | 50 | 0.5585 | 36.1231 | 1 |
| SKW-1_smr2016 | 50 | 0.9313 | 98.302 | 1 |
| SKW-1_spr2014 | 50 | 0.9862 | 23.9365 | 1 |
| SKW-1_spr2015 | 50 | 0.9394 | 2.79132 | 1 |
| SKW-2_smr2014 | 50 | 0.8141 | 68.205 | 1 |
| SKW-2_smr2015 | 43 | 0.8222 | 72.0856 | 1 |
| SKW-2_smr2017 | 50 | 0.6544 | 50.0446 | 1 |
| SKW-2_smr2018 | 50 | 0.8494 | 43.5504 | 1 |
| SKW-2_spr2015 | 32 | 0.8975 | 77.6275 | 1 |
| SKW-2_spr2018 | 50 | 0.8554 | 82.2986 | 1 |

**Table S2: Extinction–recolonization events as inferred from field observations.** True positives according to genomics data are in bold. False positives (non-bold) might represent situations where animals remained undetected or were only present as resting stages.

| Sample before | Sample after | Median MAF before | Median MAF after | Pairwise *F*_ST_ | Total change, gain, and loss of SNPs |
| --- | --- | --- | --- | --- | --- |
| FS-27_spr2016 | FS-27_smr2018 | 0.24271845 | 0.26086957 | 0.07071232 | 136123,16972,-3344 |
| FS-31_smr2014 | FS-31_smr2017 | 0.22580645 | 0.22 | 0.0536393 | 313090,17583,-23349 |
| FS-3_spr2016 | **FS-3_smr2018** | **0.23076923** | **0.33333333** | **0.19738916** | **243090,3084,-104400** |
| LA-29_spr2014 | **LA-29_smr2018** | **0.27777778** | **0.35294118** | **0.49182242** | **193423,50736,-38826** |
| LON-6_smr2016 | **LON-6_smr2018** | **0.37777778** | **0.38888889** | **0.27331328** | **355617,50907,-83494** |
| LON-9A_spr2014 | **LON-9A_smr2018** | **0.29411765** | **0.35** | **0.13538736** | **201458,5417,-65700** |
| N-27_spr2016 | **N-27_smr2018** | **0.24324324** | **0.38709677** | **0.28999761** | **274605,14746,-109911** |
| N-28A_smr2016 | **N-28A_smr2018** | **0.39622642** | **0.38297872** | **0.3857772** | **496469,93658,-95297** |
| N-28_smr2016 | **N-28_smr2018** | **0.39130435** | **0.38095238** | **0.3857772** | **496469,93658,-95297** |
| N-71_spr2016 | N-71_spr2018 | 0.26190476 | 0.19642857 | 0.32354326 | 407565,259996,-8335 |
